# The dengue virus NS1 protein alters *Aedes aegypti* midgut permeability and favors virus dissemination

**DOI:** 10.1101/2025.07.15.663935

**Authors:** Quezada-Ruiz Edgar, Silva-Olivares Angélica, Talamás-Lara Daniel, Diaz Grecia, Alanís-Muñoz Sofia, Cruz Raymundo, Betanzos Abigail, Hernández-Martínez Salvador, Lanz-Mendoza Humberto, Juan E. Ludert

## Abstract

The dengue virus is an arbovirus of public health importance transmitted by *Aedes aegypti* mosquitoes. The NS1 protein is a multifunctional glycoprotein, highly conserved among Orthoflaviviruses, that is secreted from infected cells, circulate in the sera of dengue patients at high concentrations and has been involved with pathogenesis by several mechanisms, including vascular leakage and immune evasion. However, the role of NS1 within the mosquito vector is not fully understood. In this study, we observed that feeding female *Aedes aegypti* mosquitoes with blood meals supplemented with recombinant NS1 resulted in increased midgut permeability, as evaluated by the diffusion of the non-toxic dye Brilliant Blue FCF into the hemocoel. Histological and transmission electron microscopy analysis revealed epithelial damage and disruption of the midgut cellular junctions. Immunofluorescence assays further demonstrated the delocalization of septate junction proteins essential for epithelial integrity. In addition, metalloproteinase expression in the midgut was also reduced. All these effects were abolished when NS1 was heat-inactivated indicating that they were NS1-dependent. Significantly, virus blood meals containing NS1 resulted in enhanced dissemination of DENV to secondary organs and earlier virus presence in the mosquito salivary glands, in comparison with meals treated with specific NS1 antibodies. Our findings reveal a novel function for NS1 in mosquitoes, and expand the understanding of NS1 functions beyond its human pathophysiological role. In addition, highlight NS1 as a strategic intervention point to reduce transmission efficiency in mosquitoes.

**Importance:** Dengue is the most important virus disease transmitted by mosquitoes to humans. Thus, a deeper understanding of the virus-vector interaction is required for the development of control measures and the mosquito transmission capacity. In this work, we present evidence indicating that the NS1 protein plays a role in the establishment and dissemination of the dengue virus infection in *Aedes aegypti* mosquitoes. Ingested NS1 was found to disrupt the septate junctions and to inhibit metalloproteinase expression of the midgut, allowing the virus to escape the digestive tract. In addition, NS1 was found to promote virus dissemination into the hemocoel, carcass and salivary glands. These findings uncover new functions for NS1 in the mosquito and highpoint ways to interfere with vector competence.

## Introduction

Dengue fever is the most frequent mosquito borne viral disease affecting humans, and poses a significant socioeconomic burden, particularly in tropical and subtropical regions of the planet (1). Infection may be asymptomatic, or present clinical symptoms ranging from a mild fever to high fever, headache, retro-orbital pain, muscle and joint pain, and skin rash. In some cases, dengue fever can progress to severe dengue, characterized by respiratory distress, plasma leakage, severe bleeding and organ dysfunction (2). Each year, an estimated 390 million infections occur globally, of which approximately 100 million are symptomatic, and 25,000 results in death (3).

The dengue virus (DENV) is an enveloped virus of approximately 50 nm in diameter, belonging to the family *Flaviviridae*, genus *Orthoflavivirus,* and exists as four antigenically distinct serotypes (DENV1-4) (2). The viral genome consists of a single-stranded, positive sense RNA approximately 10.8 kb in length, that encodes a single polyprotein that is cleaved to generate 3 structural proteins (C, M, and E) and 7 nonstructural proteins (NS1, NS2A, NS2B, NS3, NS4A, NS4B, NS5) (4). The non-structural protein 1 (NS1) is a multifunctional protein that is in the lumen of the endoplasmic reticulum, part of the viral replication complexes, but is also secreted along with the virions (5). Circulating NS1 has been linked to pathogenesis by several mechanisms, including complement fixation, generation of host cross-reactive antibodies, induction of pro-inflammatory cytokines and alteration of vascular homeostasis (6–8). Actually, the circulating NS1 protein of several orthoflavivirus increases the expression and release of heparanases and sialidases by endothelial cells. When these enzymes are released, they cause degradation of the endothelial glycocalyx, which may contribute to the severe manifestations of the disease, such as plasma leakage (9,10). In addition, it has been reported that the NS1 of DENV causes the delocalization and phosphorylation of endothelial cell tight junction proteins, such as ZO-1, β-catenin, and VE-cadherin (11) thereby increasing vascular permeability.

DENV are transmitted primarily by *Aedes aegypti* and *Aedes albopictus* (12). When feeding on an infected host, the mosquito acquires the NS1 protein in conjunction with the virion. In the mosquito midgut, the virus undergoes replication and subsequently disseminates to secondary tissues, such as fat body and hemolymph, finally reaching the salivary glands (13). This replicative process in the mosquito is known as extrinsic incubation period, and typically lasts 8 to12 days (14,15). In the mosquito digestive tract, essential physiological events such as protein and nutrient absorption, occur in the midgut (15), which is composed of a single monolayer of epithelial cells and a basal lamina (15,16). While absorption is carried out by the epithelial cells, the basal lamina, composed mainly of laminin and collagen IV constitute an impermeable barrier so tight that they prevent the passage of particles larger than 10 nm (17,18). It is known that the distension of the midgut due to blood feeding causes tissue damage (19, 21) and microfractures in the cell monolayer and basal lamina that create escape routes for viruses towards the hemocoel (19, 22). Also, activation of metalloproteinases and other enzymes involved in repairing the basal lamina by pathogen exposure may result in weakening of the lamina (23). However, the mechanisms used by DENV and other orthoflaviviruses to escape the midgut are not fully understood. It has been shown that the DENV NS1 protein can downregulate the transcription of key immune response genes in the mosquito, potentially facilitating infection in the midgut (24). However, it is unknonw if the reported ability of NS1 to permeabilize host endothelia also occur in the intestinal epithelium of the vector. In this study, we investigate the role of DENV NS1 in altering *Aedes aegypti* midgut permeability and its role in virus dissemination and implications for vectorial competence.

## Materials and Methods

### Mosquito rearing

Adult *Aedes aegypti* Rockefeller strain mosquitoes were reared in the insectary of Instituto Nacional de Salud Publica (INSP), Mexico, under the following conditions: 60-80% humidity, 28-30°C, and a 12:12h light-dark cycle. The larvae were fed with tropical fish food (*TetraMin* flake fish food). The adult stage was maintained in cages, with ad libitum access to a 10% sugar solution. Adult females of 5 to 7 days of age were used for subsequent experiments.

### Organ dissection

Mosquitoes were coldly anesthetized and transferred to slides with a drop of Phosphate-Buffered Saline (NaCl 137 mM, KCl 2.7 mM, Na_2_HPO_4_ 10 mM, KH_2_PO_4_ 1.8 mM). Dissection was performed using a stereo microscope.

#### Midgut

A transverse cut was made in the head and the antepenultimate abdominal segment. The mosquito’s body was separated from the abdomen by holding it with forceps, and the midgut was removed intact. Tissues were washed with PBS until the ingested blood was removed, and then they were placed in microcentrifuge tubes containing 50 µl of PBS.

#### Carcass

Carcasses were recovered after removing the midguts as described above and placed in microcentrifuge tubes containing 50 µl of PBS. Tissues were stored at −70°C until used.

### Cell cultures and virus propagation

Baby hamster kidney cells (BHK-21, ATCC CCL-10) were cultured on DMEM medium supplemented with 5% heat-inactivated fetal bovine serum (FBS, Gibco) under standard conditions of 37 °C with 5% CO2. DENV-2 strain New Guinea C (NGC) was propagated in BHK cells using neonatal mouse brain extract with an MOI of 0.1 in DMEM medium without FBS. After 5 days post-infection, the supernatants were collected and clarified by centrifugation at 12,000 *g* for 15 min at 4°C.

The virus was quantified by two-step qPCR using the reagent Maxima SYBR Green/ROX qPCR MasterMix (2x) (Cat. no. K0221) on a Rotor-Gene 5Q (Qiagen). 800ng of total RNA was used as a template. For cDNA generation, 1 µl of random hexamer primer (0.2 µg/µl; Cat. no. SO142), 1 µl of dNTPs (10 mM; Cat. no. R0192) and 1 µl of RevertAid (200 U/µl; Cat. no. EP0441) were used in a final volume of 20 µl. DENV primers were used for the mix at a final concentration of 0.3 µM. The reaction parameters were as follows: 10 minutes at 95°C, and a 40-cycle PCR reaction with the following settings: 95°C for 20 seconds and 60°C for 45 seconds. A standard curve using a 10-fold serial dilution of a synthetic gene containing the primer targets flanking regions (gBlock) was used for absolute quantification (25). The Ct values obtained were extrapolated from a standard curve created using the performed dilutions. The concentration obtained was approximately 1×10^7^ viral RNA copies/ml. The sequence of the primers and the gblock can be found in Supplementary Table 1.

### Membrane blood feeding

Adult female mosquitoes were deprived of sugar for 12 hours to facilitate treatment administration. Mosquitoes were artificially fed with heparinized rabbit blood at 250 µl/L in combination with DMEM medium in a final volume of 1 ml for each treatment. Experimental conditions were as follows: i) Negative, untreated control: Blood + DMEM medium 1:1 (v/v). ii) Positive, Dithiothreitol (DTT)-treated control: 2.5 mM DTT final concentration. iii) Pure recombinant NS1: One µg recombinant NS1 DENV-2 (rNS1) (Biotechne, Cat. no. 9439-DG-100), and iv) Pure recombinant NS1 heat-denatured: One µg rNS1 boiled in a water bath for 30 min.

### Intestinal permeability assays

A modified version of the ‘Smurf assay’ (26, 27) was used to assess changes in midgut epithelial permeability. Brilliant Blue FCF (Merck, Cat. no. 80717) was prepared at 1% in PBS. Twelve hours before the assay, female mosquitoes were fed a 1% dye-supplemented sucrose solution. On the following day, they were starved for 6 hours and then fed with rabbit blood mixed with the dye. Hemolymph was collected from individuals exhibiting a blue coloration in their legs, indicating dye leakage into the hemocoel. PBS was injected into the thorax region using a microinjector (Nanoject I - Drummond Scientific). Subsequently, we recovered the traces of dye that reached the coelomic cavity by making an incision in the last segment of the abdomen and collecting the hemolymph with a micropipette. The hemolymph was clarified by centrifugation at 12,000xg for 10 minutes to remove hemocytes and fat bodies, and absorbance was measured at a wavelength of 630 nm using a spectrometer (Elisa Plate Reader, DASITALY). The reducing, permeabilizing agent DTT was used as a and positive control (18). Values were expressed in relative units. Dye outflow was assessed at 12, 24, and 36 h after dye feeding or post-treatment (hpt). Three biologically independent experiments were performed, and triplicate samples were collected for each treatment.

### Hematoxylin-eosin stains

Twenty-four hours after administration of the treatments, abdomens from fully fed adult females were dissected, fixed in Bouin’s solution (5% acetic acid, 9% formaldehyde, 0.9% picric acid (Sigma-Aldrich) for 10 minutes at 56°C and kept overnight at room temperature (RT). Next day samples were transferred to 70% ethanol before embedding in paraffin. Samples were dehydrated in an ethanol-xylene lane (50%, 70%, 80% 95%, 100%, Xylene) and embedded in Paraplast X-TRA (Oxford, St. Louis, MO) overnight at 56°C (Celestino-Montes et al. 2021). Abdomens were mounted in Paraplast X-Tra using histological cassettes (Quebec, Canada). Six µm-thick histological sections were made in a longitudinal plane and mounted on slides pre-treated with 1% gelatin (Sigma) (28). Finally, slides were hydrated and stained using the standard Harris hematoxylin-eosin protocol (Harris hematoxylin (Sigma Aldrich, St. Louis, MO, USA). Five female mosquitoes’ abdomens per treatment were examined with three biological replicates.

### Transmission electron microscopy

#### Sample preparation and paracellular permeability assay

Five midguts per treatment were dissected as described above, fixed with 2.5% glutaraldehyde dissolved in 0.1 M sodium cacodylate buffer, pH 7.2 at RT, and post-fixed with 1% osmium tetroxide. After dehydration in increasing concentrations of alcohol, the samples were pre-embedded in a mixture of 100% alcohol and Polybed resin (2:1 and 1:1), before embedding in pure resin. The resin was left to polymerize at 60 °C for 24 hours. Subsequently, 60 nm thin sections were made and contrasted with uranyl acetate and lead citrate. To visualize the septate junctions, samples were further stained with 0.6% ruthenium red, as described (29, 30) and finally analyzed with a JEOL-JEM-1400 transmission electron microscope (JEOL Ltd., Tokyo, Japan).

### Localization septate junction proteins

#### Sample preparation

To evaluate changes in the localization of proteins involved in the septate junctions, female abdomens collected 24 hours post-treatment, were dissected as described above and fixed for 2 hours at RT in 4% paraformaldehyde in PBS, pH 7.2. After three washes with PBS, the abdomens were placed in 10% sucrose overnight. Samples were embedded in tissue freezing medium (Leica Microsystems, Wetzlar, DE) and immediately frozen at −20 ^°^C.

#### Immunofluorescence

The mounted abdomens were cryosectioned in the transversal plane of the tissue (cryostat Leica CM1100, Leica Microsystems), making 6 µm thick sections cuts only in the midgut section, adhering them on slides pretreated with 0.5% gelatin (Sigma) and potassium dichromate (Merck) 0.05%. Sections were permeabilized with methanol at −20°C for 7 min in cold, followed by blocking with 1% BSA in PBS 0.1% Tween-20. Rabbit anti-claudin-1 (1:100; Invitrogen, cat. no. 51900) and mouse anti-β-catenin (1:50; Santa Cruz cat. no. sc376959) used as primary antibodies were incubated with the sections at 4°C overnight. After three washes with PBS, sections were incubated with goat anti-rabbit and mouse anti-IgG conjugated with AlexaFluor 488 and AlexaFluor 647, respectively (Invitrogen), for 2h at RT. Nuclei were counter stained with DAPI solution (1:800; Sigma D9542) for 10 minutes. Each slice was mounted in 5 µl of VectaShield (Vector, H-1000) and analyzed on an LSM 900 confocal microscope. Images were processed with Zeiss Zen Blue 3.6 software. Three abdomens of female mosquitoes per treatment were examined.

### Dissemination of DENV to secondary tissues

To evaluate the effect of NS1 on virus dissemination, mosquitoes were fed with DENV-2 from BHK cells incubated for 1h at RT either with purified rabbit pre-immune serum at a concentration of 8µg of IgG (1:100), or purified rabbit polyclonal anti-NS1 at a concentration of 9µg of IgG (1:100). Infection was evaluated at 24, 48, and 72h post feeding in midgut and carcass. In addition, mosquito’s saliva was extracted and analyzed 7 dpi (31). Total RNA was extracted and retro transcribed as described above. PCR were performed using 100 ng of cDNA as a template. The primers used are described in Supplementary Table 1. Relative quantification was performed using the endogenous gene *eEF1*α with the Pfaffl method (32). The infection ratio was calculated by counting all positive tissues among the total number of tissues evaluated at each time point, and results expressed as percentage. Three biological replicates were made collecting 5 pools of 5 tissues per replicate.

### Statistical analysis

A normality test was performed for each data before employing any statistical test. For data with a normal distribution, a One-way ANOVA Tukey test was used, and for not-normally distributed data a nonparametric test (One-way ANOVA/ Kruskall-Wallis test) w. Infection prevalence was compared using a Fisher’s exact test. A 2-way ANOVA/Bonferroni’s multiple comparisons test was used to compare the spread of the virus to secondary tissues. The criteria used were *p<0.5, **p<0.05, ***p<0.001. All analyses were performed and graphs using GraphPad Prism 8.0.2. statistical software (33).

## Results

### NS1 protein increases midgut epithelial permeability in *Aedes aegypti*

To determine if the NS1 protein would affect midgut permeability, the outflow of blue dye No. 1 was quantified in mosquitoes fed with rNS1. Initially, a dose-response assay was performed using increasing concentrations (0.062 to 1.0 µg) of rNS1, and collecting the samples 24 hpt (Fig 1a). Although a plateau was not reached, the results showed a clear dose-dependent effect, with the 1 µg/ml concentration causing the most significant dye leakage into the hemocoel (Fig 1b). Next, using this rNS1 concentration a time course at 12, 24, and 36hpt of dye outflow was carried out, together with heat inactivated rNS1 and an untreated condition as negative controls, and DTT as positive control. Mosquitoes fed with DTT showed a significant outflow of the dye at 12 and 24hpt (Fig.1d-e), indicating disturbance in the midgut epithelium, presumably due disruption of the disulfide bonds that stabilize the proteins conforming the septate junctions (18). Interestingly, rNS1 induced changes at 24 hpt similar to the ones observed with DTT (C-; p<0.01), and that were not observed in the heat denatured condition (p<.05). The effects caused by the rNS1 were not observed at 12 hpt, suggesting a delay in relation to DTT induced changes. Finally, at 36 hpt, no dye outflow with any of the treatments, even DTT, were observed, suggesting that the disruption effect was reversed (Fig. 1f). All these results indicate that NS1 can cause midgut barrier dysfunction.

**Figure 1.**
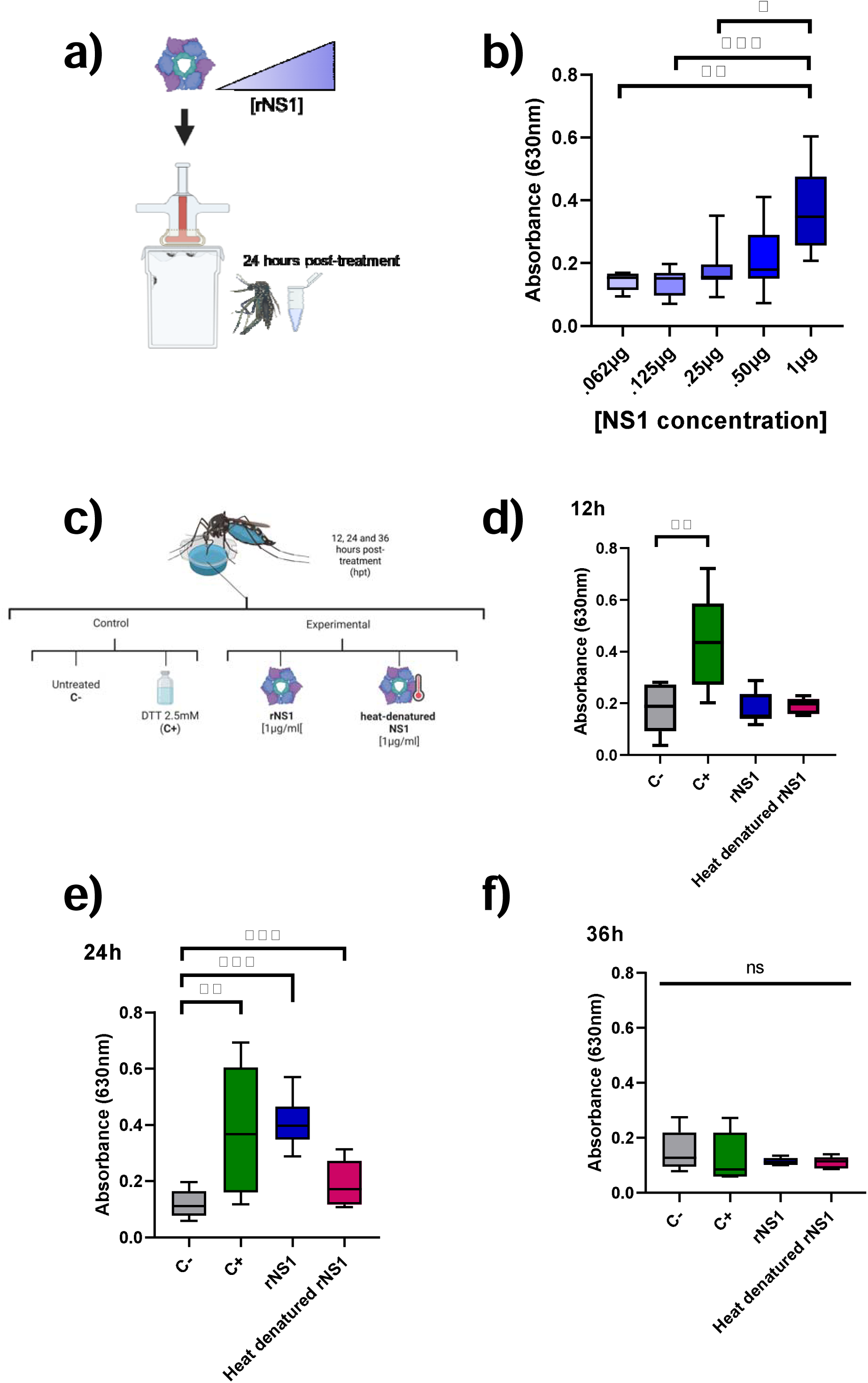
The presence of recombinant NS1 (rNS1) favors the transient outflow of the dye into the hemocoel. (a) Schematic representation of the experimental design for the dose-response assay with rNS1. (b) Dose-dependent increase of midgut permeability observed in response to rNS1: OD measure of Brilliant Blue FCF presence in haemolymph. (c) Experimental conditions used to evaluate the role of native and denatured NS1 in modulating epithelial permeability, including untreated (C−), Dithiothreitol 2.5mM (C+), recombinant NS1 (rNS1), and heat-denatured rNS1. Quantification of dye outflow at 12 (d), 24 (e), and 36 (f) hours post-treatment (hpt). rNS1 significantly increased permeability compared to controls at 24 hpt, while heat-denatured rNS1 did not induce a significant effect. No dye leakage was observed at 12 and 36 hpt. Three biological replicates per assay were performed (n= 90). ANOVA/Tukey *p<0.05 **<0.01 ***<.001.

### rNS1 protein causes a histological alteration in the midgut

Histological analysis of the midgut of the mosquitoes exposed to rNS1 was performed on longitudinal sections of the abdomen. Previously, in other to stablish the half-life of the feed rNS1, or how long it will remain in the midgut, dot-blot assays were carried out. NS1 was observed in the midguts at 24, but not at 48, hpt, suggesting exit or degradation of NS1 (Supp. Fig. 1). Thus, the following experiments were all performed at 24 hpt.

Hematoxylin-eosin staining showed a loss in the microvilli of epithelial cells in the midguts of the mosquitoes fed with blood and rNS1 protein when compared to the control, untreated condition (Fig. 2a-c). This effect was not observed when heat denatured rNS1 was used (Fig. 2d). No evident morphological alterations were observed in the midgut epithelium of mosquitoes treated with DTT, except some loss of mucus continuity when compared to untreated controls (Fig. 2b).

**Figure 2.**
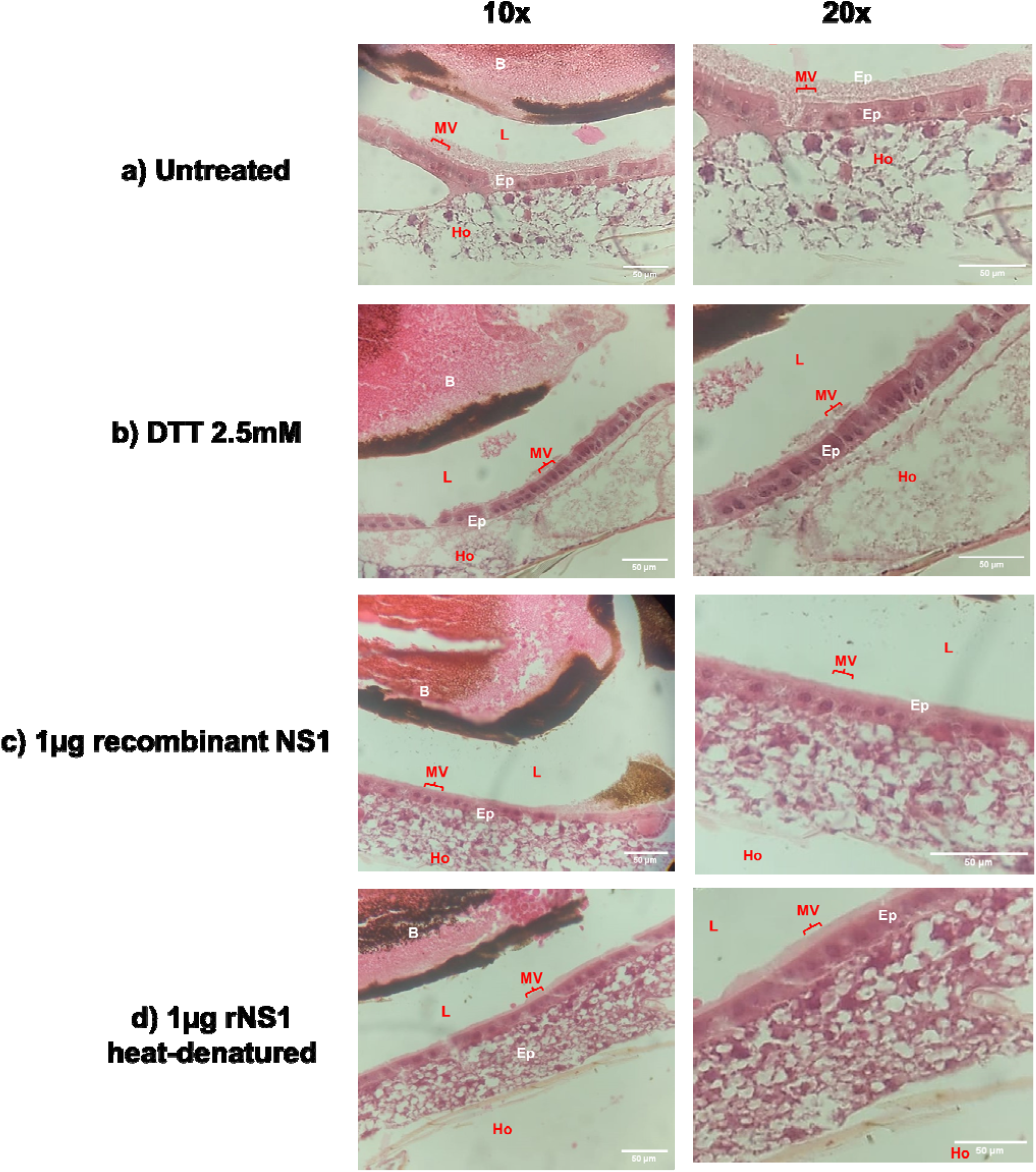
Microvilli reduction in midguts of *Aedes aegypti* exposed to rNS1 protein. Longitudinal views of H-E-stained sections of the epithelial midgut. Structural changes were observed in the microvilli zone when rNS1 was present (c). There was no reduction in microvilli when rNS1 was heat denatured (d) absent (a) or treated with DTT (b). Three abdomens of female mosquitoes per treatment were examined with reproducible results. Lumen (L), epithelial cells (Ep), microvilli (MV), and hemocoel (Ho). Scale bar: 50 μm.

The losses in cell microvilli were confirmed by TEM. Ultrastructural analysis of the mosquito midgut showed reduction in the number of microvilli in the rNS1-fed mosquitoes (Figure 3b), in contrast with the untreated condition (Figure 3a), or mosquito treated with heat denatured rNS1 (Figure 3c), which showed no changes and maintained dense apical microvilli. No morphological changes were evident in the cell nuclei (Fig 3d-f) or in the basal lamina of the examined midgut epithelial cells (Fig. 3g-i).

**Figure 3.**
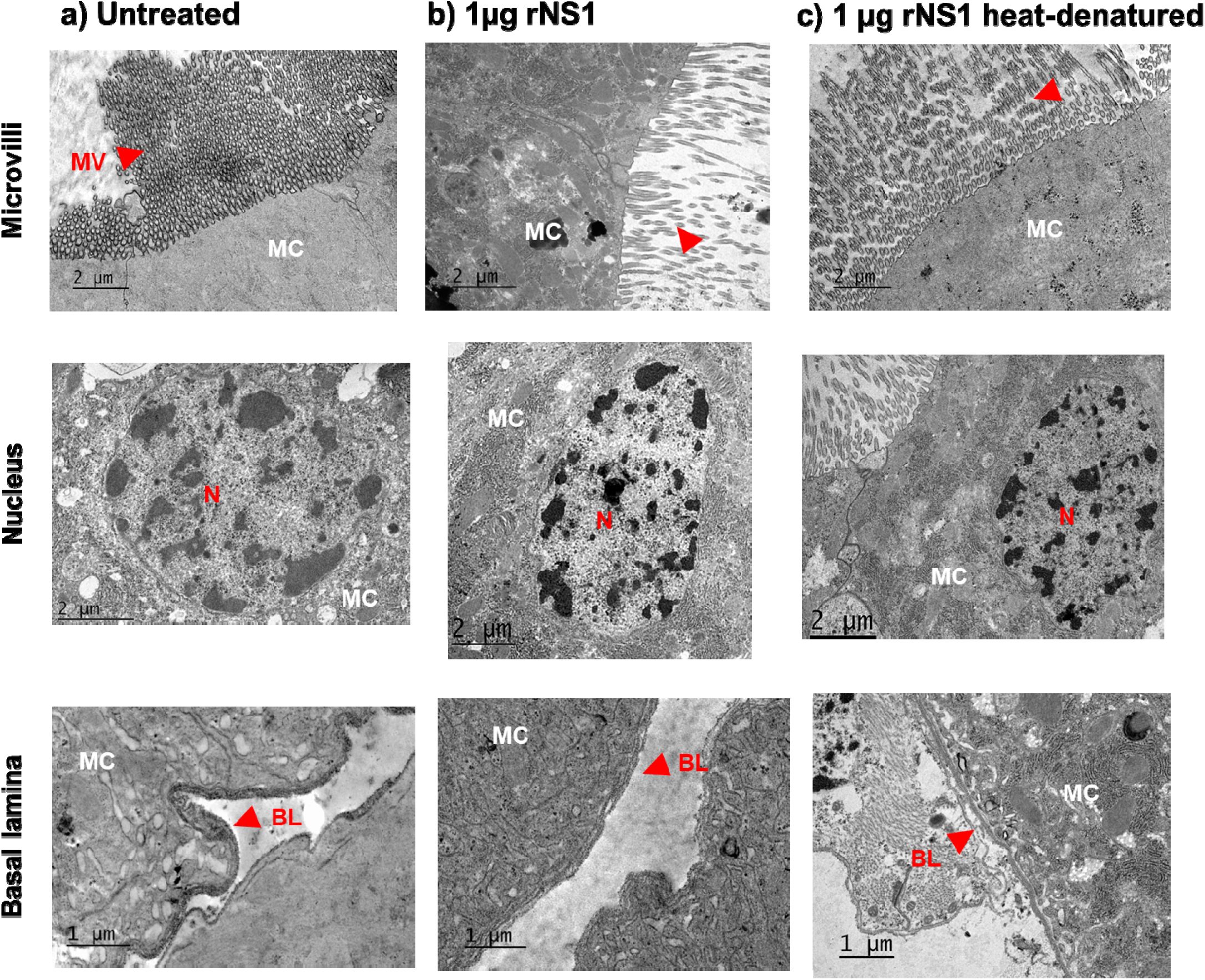
Transmission electron microscopy (TEM) analysis of the midgut epithelial cells of *Aedes aegypti* exposed to rNS1 protein. The red arrows indicate the microvilli zone. Microvilli in the apical zone of the midgut epithelial cells of the untreated group were numerous and appear undamaged (a); in the presence of the rNS1 protein there appeared to be a reduction in the number and length of the microvilli (b), a damage that was reversed when the protein was denatured by heat (c). No other changes were observed in cell morphology, nor the shape of the nucleus and basal lamina. MC: midgut epithelial cells, MV: microvilli, N: nucleus, BL: basal lamina. Five midguts of female mosquitoes per treatment were examined with reproducible results.

### The NS1 protein induces changes in the paracellular permeability of the *Aedes aegypti* midgut epithelium

To further investigate the midgut changes caused by rNS1 protein, cell-to-cell junctions and septate junctions (SJ) integrity was evaluated using ruthenium red as an indicator of altered paracellular permeability. Alteration indicators include mainly how far the ruthenium red is incorporated and the width of the septa, but also morphological or perimeter and area variations in the spots located at the septate junctions or width of the areas where ruthenium red is not intercalated (Supp. Figs. 2 and 3). Ruthenium red intercalation at the cell junctions was observed in all conditions (Fig. 4a-c). However, the length of dye penetration was significantly longer in the rNS1 compared with the untreated midguts or treated with heat-denatured protein (Fig. 4d). Channel width was also larger in the samples with rNS1 than in controls, albeit the width of these channels was even greater in the NS1-heat denatured condition than in the other conditions (Fig. 4e).

**Figure 4.**
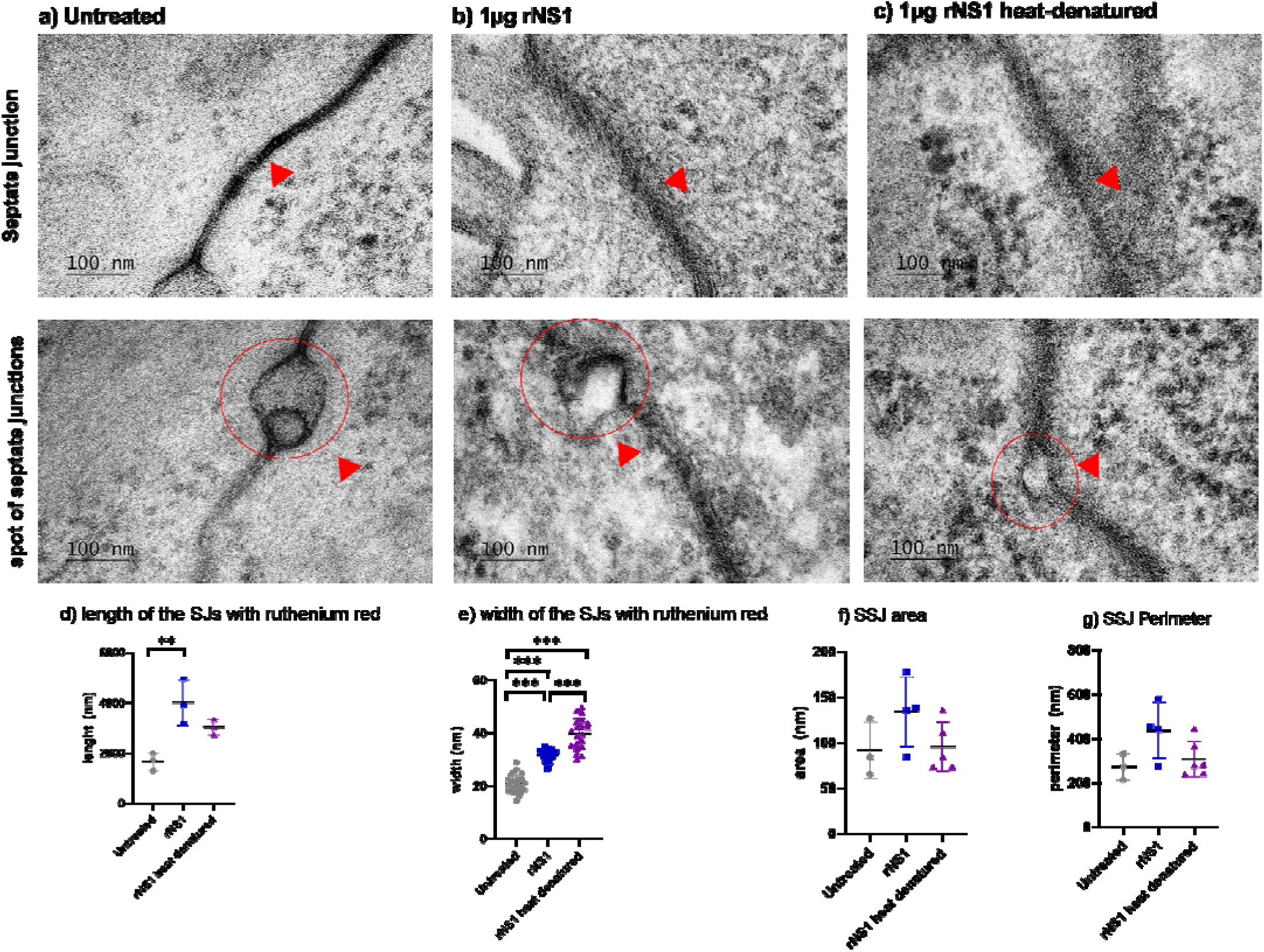
Qualitative changes in the septate junctions (SJ) and spot of septate junctions (SSJ) of *Aedes aegypti* exposed to rNS1 protein. The spot is indicated by surrounding it with a red circle and pointing at it with red arrows. In untreated mosquitoes (a), the ruthenium red (red arrow) is incorporated into the SJ but is kept into the septum. The SSJ remained intact. When the mosquitoes ingested rNS1, ruthenium red emerged from the SJ septum, SSJs lost their shape (b). Denaturation by heat of the rNS1 did not prevent ruthenium red from leaving the septum but no changes in the SSJ were observed (c). Measurements were made using Image J. The highest ruthenium red incorporation was observed in the presence of rNS1 (d). The width of the septa when ruthenium red was incorporated (e). No changes were observed in areas (f) and perimeter (g). Images from three independent experiments were analyzed using Fiji/Image J software. The * above icons indicate significant differences between conditions. ANOVA/Tukey test *p<0.05 **<0.01 ***<.001. Scale bar: 100 nm.

Sac-like structures, called spots septate junctions (SSJs), characterized as intercellular cavities or gaps between cell-cell junctions were also examined (34, 35). There was a loss of integrity of these spots in the presence of rNS1 protein compared to untreated midguts (Fig. 4a-b), and their morphology was unaffected when NS1 was denatured (Fig. 4c). None of the treatments significantly changed the area or the perimeter of the SSJs (Fig. 4d-e). No changes were observed in the regions of the SJs where ruthenium red was not intercalated (Supp. Fig. 3a-c); however, the width of the septa was altered, being widest in the presence of rNS1 protein (Supp. Fig. 3d). These ultrastructural alterations taken together support the idea that rNS1 does change the integrity of the epithelial barrier in mosquito midguts through remodeling of septate junctions.

### NS1 protein causes delocalization of proteins conforming the midgut septate junctions

The integrity of the septate junctions was also analyzed by confocal immunofluorescence microscopy. However, previously an in silico comparative analysis using available protein databases was carried out to find potential *Aedes aegypti* proteins involved in the formation and maintenance of septate junctions. Two candidate proteins were chosen based on their similarity to known cell junction complex components in other organisms: AAEL001447 (TMEM47), a claudin-like transmembrane protein, and AAEL002887 (ARM), an orthologue of the Armadillo segment polarity protein. Both proteins are related to cell-cell junction proteins found in vertebrates (claudins and β-catenins, respectively). Phylogenetic analysis and domain prediction confirmed the relationship with these types of protein (Supp. Fig. 4). Thus, these 2 candidate proteins present domains likely to be labelled with antibodies directed to claudin-1 and β-catenin of vertebrates. Therefore, we analyzed the localization of these candidates in the mosquito midguts exposed to rNS1 protein.

In an unaltered, control midgut, TMEM47 is strictly localized in the apical region and ARM in the basal region of the epithelial cell (Fig. 5a). The presence of the rNS1 protein provokes a delocalization of TMEM47 from the apical to the basal region of the epithelial cells. Interestingly, the localization of the ARM protein also changed, delocalizing mainly to cytoplasm and nuclei (Fig. 5b). None of these changes were observed when the rNS1 protein was denatured (Fig. 5c). These results indicate that NS1 alters the location of key component proteins of SJs.

**Figure 5.**
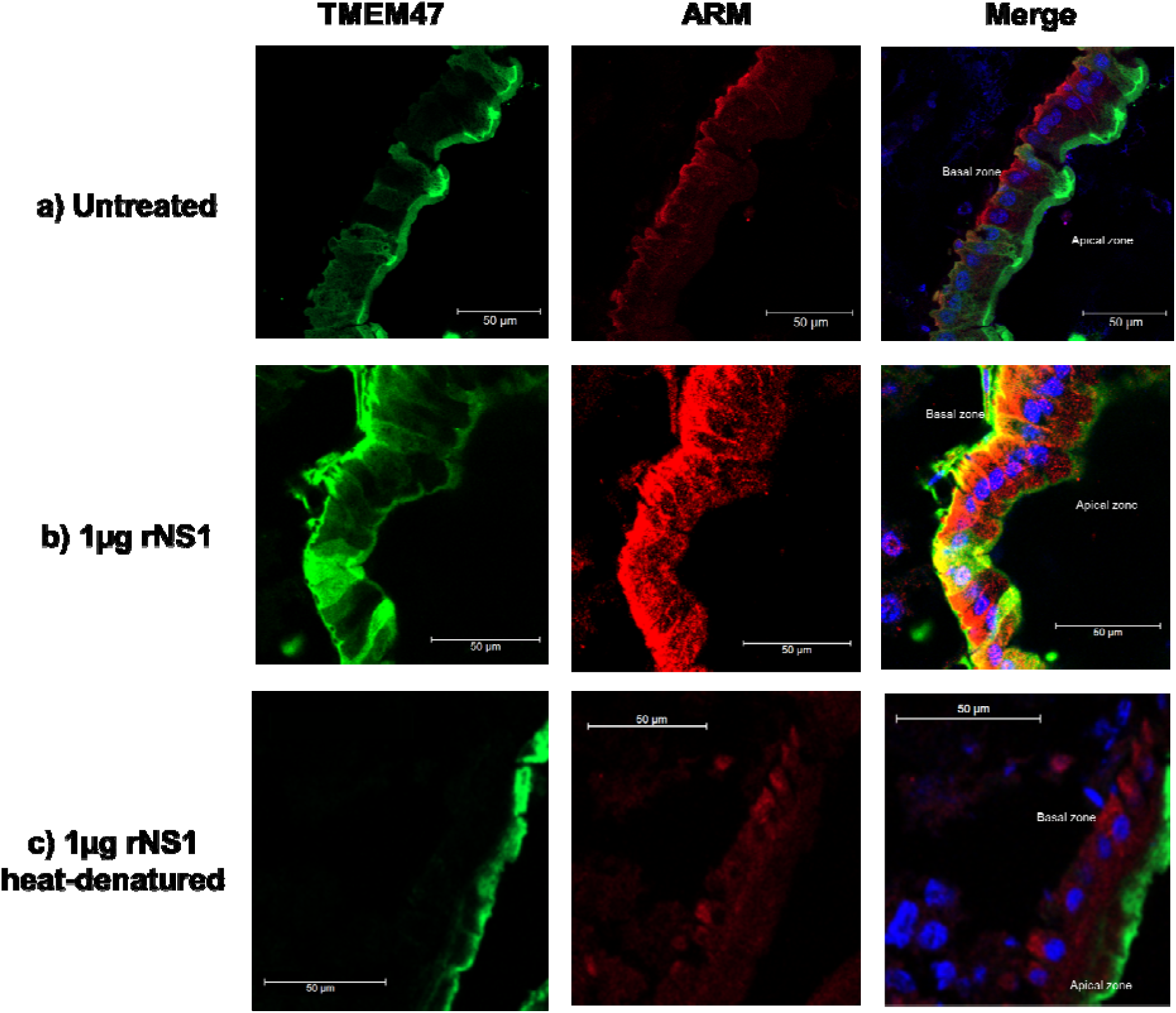
Confocal immunofluorescence microscopy analysis of TMEM47 (green) and ARM (red) proteins in transverse cryosections of *Aedes aegypti* midguts. (a) In untreated midguts, TMEM47 localizes at the apical region of the epithelial cells while ARM is present at the basal region. (b) rNS1 treatment leads to TMEM47 relocalization toward both apical and basal compartments, whereas ARM is primarily redistributed to the cytoplasmic compartment and nuclei. (c) Denatured rNS1 does not alter the typical localization of either TMEM47 or ARM. Nuclei were counterstained with DAPI (blue). Three abdomens of female mosquitoes per treatment were examined with reproducible results. Scale bar: 50 μm. n≥3.

It has been reported that the alteration in tight junction proteins by NS1 is associated with changes in the expression of several metalloproteinases (36–38). Thus, the expression levels two metalloproteinases genes, Aemmp1 and Aemmp2, involved in basal lamina degradation in mosquitoes CHIKV infection (23, 39, 40), was evaluated by qRT-PCR in midguts exposed to rNS1. A significant decrease in Aemmp1 expression was observed in mosquitoes with rNS1 at 24 h post-feeding (Supp. Fig. 5), that is partially recovered if rNS1 is heat denatured. For Aemmp2, a decrease in messenger expression was observed in the presence of DENV rNS1, either heat denatured or not, at 24 and 48 hpf. However, expression is recovered at 48 hpf with the heat denatured NS1 (Supp. Fig. 5). These results suggest that inhibition of MMP expression occurs along with the delocalization of TMEM47 and ARM proteins.

### The NS1 protein promotes dengue virus dissemination

Given that DENV-2 NS1 alters the midgut epithelial barrier by increasing paracellular permeability and disrupts septate junction proteins, we hypothesized that these alterations may facilitate viral dissemination to secondary tissues. To evaluate the role of NS1 in DENV-2 dissemination, we compared viral spread in mosquitoes where NS1 function was either intact or inhibited. For this, supernatants collected from DENV infected BHK-21 cells, were incubated with pre-immune serum or hyperimmune anti-NS1 serum (1:100 for both conditions), before being fed to *Aedes aegypti* females. The capacity of these supernatants to cause midgut permeability was tested in intestine permeability assays (Fig. 6f). In addition, the supernatants were tested for the presence of NS1 using a commercial ELISA (data not shown) and dot-blot assays (Supp. Fig. 1), and virus load by qRT-PCR (data not shown). Viral dissemination was assessed in the midgut, carcass, and saliva across infection time points by qRT-PCR.

**Figure 6.**
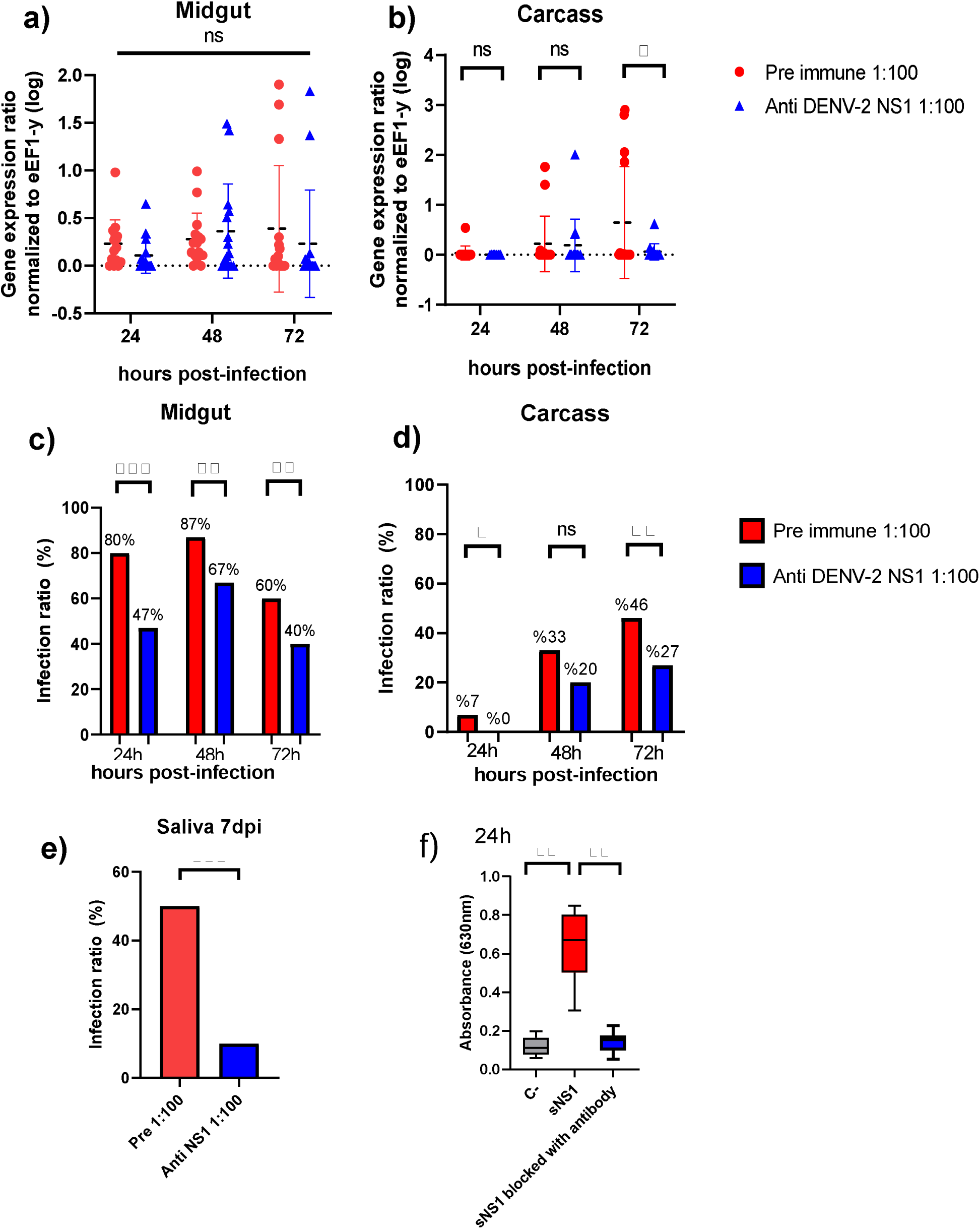
NS1 protein promotes virus dissemination to secondary tissues in *Ae. aegypti*. Female mosquitoes were orally infected with DENV-2 in combination with pre-immune serum (red) or anti-NS1 antibodies (blue). Relative quantification of viral genomes was evaluated at 24, 48, and 72 hours post-infection in midguts (a) and carcasses (b). The infective ratio was evaluated at 24, 48 and 72 hours-post infection in midguts (c) and carcasses (d). Virus presence was evaluated in salivary gland at 7 days post-infection (e). Quantification of dye outflow at 24 hours post-treatment in mosquitoes exposed to infected cell supernatants treated with pre- or hyperimmune sera to NS1. Untreated control condition same as shown in Figure 1 at 24 hpi (e). *(p < 0.05), ** (p < 0.01), ***(p < 0.001). n≥3.

Relative viral load analysis revealed no significant differences between treatments in the midgut (Fig. 6a) or carcass (Fig. 6b), except at 72 hpi in carcasses, where mosquitoes fed with unblocked NS1 exhibited higher viral loads (p < 0.05). Nontheless, infection ratios in the midgut showed significant differences starting at 24 hpi, with the unblocked NS1 group reaching an 80% infection rate (p < 0.001). This trend persisted at 48 and 72 hpi (p < 0.01) (Fig. 6c). Similarly, in the carcass, infection in the unblocked NS1 group began increasing from 24 hpi, reaching 47% at 72 hpi, compared to only 27% in the NS1-blocked group (p < 0.05) (Fig. 6d).

These results suggest that NS1 facilitates the dissemination of the virus from the midgut to secondary tissues, likely contributing to a shortened extrinsic incubation period. To further investigate this, saliva samples were analyzed at 7 days post-infection. Mosquitoes exposed to unblocked NS1 showed a significantly higher infective ratio (46%) compared to the NS1-blocked group (27%) (p < 0.01) (Fig. 6e), indicating that NS1 promotes early viral escape from the midgut and enhances vector competence.

## Discussion

The DENV NS1 protein circulates in the blood of patients during the acute phase of disease at concentrations that reach micrograms per microliter (37, 41). Thus, it is expected to be ingested along with the virions during a mosquito bite. Previous evidence indicated that ingested NS1 favors DENV infection in the mosquito by tuning down mosquito innate immunity (24). Recently, another mechanism of orthoflavivirus infection enhancement in *A. aegypti*, albeit not NS1 related and more like a “Trojan horse” model, was reported (42). In this study, we demonstrate that NS1 also favors mosquito infection and viral dissemination by a novel mechanism, the disruption of the midgut epithelial barrier of *Aedes aegypti*. Using a dye leakage assay, we found that recombinant NS1 increases midgut permeability in a dose-dependent and transient manner. This dye-based assay has been successfully employed in other models, such as *Drosophila melanogaster* (26, 27) and *Anopheles gambiae* (43) where disruption of the midgut barrier leads to dye leakage into the hemocoel. Histological and ultrastructural analyses revealed that NS1 exposure leads to microvilli loss and morphological and functional alterations in septate junctions, and immunofluorescence assays showed that these alterations go along with the delocalization of two key junctional proteins, TMEM47 and ARM. These epithelial alterations correlated with a higher infection ratio in the midgut and secondary tissues and an accelerated presence of DENV in mosquito salivary gland. Denaturation or antibody blocking of NS1 reduced these effects, suggesting that NS1 acts as a mediator of epithelial permeability, virus dissemination and vector competence.

These findings parallel the effects of DENV NS1 on vertebrate vascular endothelium, where secreted NS1 disrupts the endothelial glycocalyx and degrades tight and adherent’s junction proteins, leading to hyperpermeability (10). In human endothelial cells, NS1 recruits host metalloproteinase 9 (MMP-9) to degrade β-catenin, ZO-1, and ZO-2, further compromising cell–cell adhesion (36–38). The present work extends this paradigm to the mosquito midgut, showing that a similar junctional disruption mechanism may underline both the vascular leak in the vertebrate host and midgut barrier dysfunction in the vector.

The morphological alterations observed by TEM in the midgut microvilli may unveil a mechanism by which the NS1 protein facilitates access to its target cells, as microvilli are known to act as physical barriers that hinder pathogen interaction with epithelial surfaces (29, 44). The reduction or disorganization of these structures, could therefore reflect an NS1-driven mechanism to weaken this first line of defense. Furthermore, the increased penetration of ruthenium red between epithelial cells suggests that NS1 disrupts septate junctions, since intact junctions normally prevent the intercalation of this tracer (29). Also, the NS1 alters the morphology of the spot septate junctions (SSJs) without changing their overall area or perimeter. The altered ultrastructure we observed supports the notion that NS1 compromises epithelial integrity, thereby enhancing paracellular permeability (34, 35).

These findings are in line with previous reports showing that ultrastructural disruptions of the basal lamina in *Ae. aegypti* are associated with increased arboviral dissemination (20, 22, 40). Secondary blood meal induces transient basal lamina damage that facilitates dengue virus escape from the midgut. Similarly, TEM studies of Chikungunya virus infection have documented localized basal lamina breaks associated with enhanced viral spread (40). These parallels suggest that NS1-induced epithelial remodeling may act in synergy with natural physiological processes or viral strategies to promote early systemic infection in the mosquito.

Our findings also revealed that rNS1 caused delocalization of key SJ components, namely TMEM47 (claudin-like) and ARM (β-catenin orthologue), associated with septate junction integrity and barrier functions in the mosquito midgut epithelium. These results are consistent with the behavior of smooth SJ in *Drosophila*, where proteins such as Mesh, Ssk, and Tsp2A are critical for barrier function and display mutual dependency for proper membrane localization. Disruption of any of these components leads to junctional breakdown, cytoplasmic mislocalization, and increased paracellular permeability (45). TMEM47, in particular, belongs to the claudin superfamily and has been proposed as a functional analog of Ssk or claudin-like proteins in mosquitoes, while ARM shares structural features with β-catenin and may be involved in anchoring cytoplasmic complexes at the junction. These changes also mirror phenotypes observed in genetic knockdown models of sSJ proteins in *Drosophila*, where similar relocalization and leakage phenotypes are reported (46). Thus, the observed altered localization patterns are therefore indicative of NS1-induced disassembly of the septate junction complex, compromising the epithelial integrity and likely facilitating paracellular viral dissemination. Despite these advances, several limitations must be considered. First, we used recombinant NS1 added externally, which may not fully reflect the dynamics or state of NS1 circulating in the sera of patients, where NS1 has been also reported to circulate complexed with HDL and LDL (47, 48). Second, only NS1 from DENV-2 was used, and albeit NS1 is a highly conserved (49), confirmation with other Orthoflavivirus NS1 is required. Third, the Rockefeller stain of *A. aegypti*, highly susceptible for DENV, was used and virus dissemination may differ in natural mosquito populations.

Midgut is the first tissue where the virus interacts with the vector (11, 50, 51). Therefore, overcoming the midgut infection barrier (MIB), is essential for viral establishment. Our results taken together indicate that NS1 i) directly contributed to the early establishment of DENV in the midgut by epithelial permeability and junctional remodeling and subversion of tissue-repair. This novel mechanical function will pair with physiological stimuli such as blood feeding, which by itself induces microfractures in the midgut epithelium, and with the immunomodulatory capacity reported for NS. Recent studies reporting that DENV and ZIKV sNS1 present in the blood meal can inhibit the JAK–STAT pathway in *Ae. aegypti* midgut cells, thereby favoring viral establishment (24, 54) It will be worth to evaluate if NS1 may also participate in other cellular processes such as apoptosis or cytoskeletal remodeling, mechanisms tightly linked to septate junction integrity, which have been explored in *the Ae. aegypti-DENV* model, but not in the context of NS1’s involvement (53–55) facilitates faster viral escape from the midgut into secondary tissues (e.g., carcass) and ultimately the saliva. In this regard, the contribution of endogenously synthetized NS1 need to be evaluated. Mosquito cells are known to secrete NS1 (56, 57) but its function may vary depending on the tissue type, such as fat body or hemocytes. Figure 7 presents a summary of the main actions of NS1 in the mosquito. Finally, these findings suggesting that NS1 reduces the extrinsic incubation period and enhances vector competence. Thus, strategies targeting NS1 could reduce the vector competence to DENV infection, offering a novel venue for dengue control.

**Figure 7.**
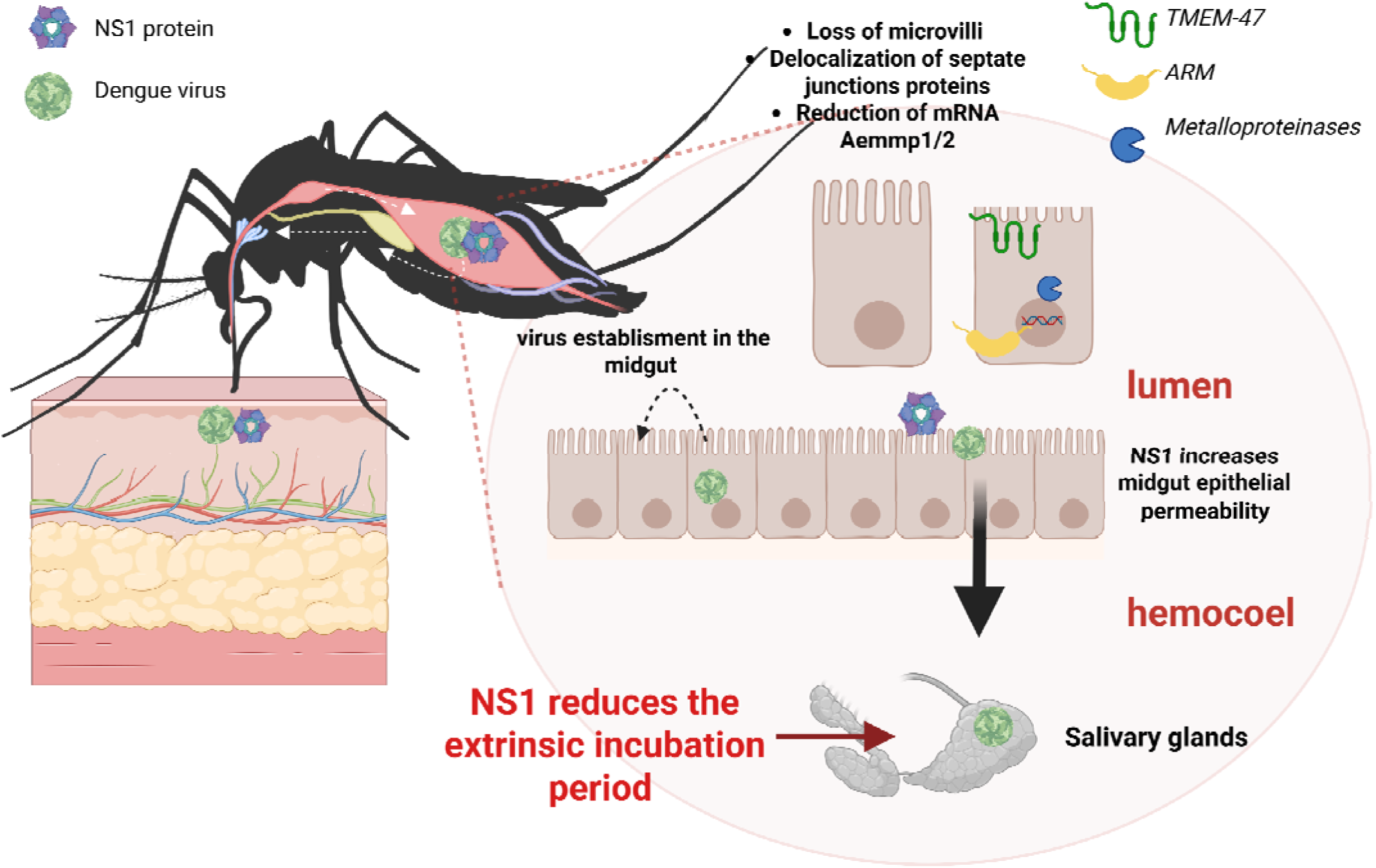
Scheme summarizing the effects of NS1 to the intestinal midgut barrier and on virus dissemination in the mosquito. Upon exposure to NS1, enterocyte show loss of microvilli, delocalization of septate junction proteins *(TMEM-47 and ARM),* and reduced expression of *AeMMP1* and *AeMMP2* genes. These alterations increase epithelial permeability, facilitating early viral escape into the hemocoel and rapid dissemination to the salivary glands. Overall, NS1 reduces the extrinsic incubation period, potentially enhancing mosquito transmission efficiency.

## Acknowledgments

We would like to acknowledge Drs. Victoria Pando-Robles, Renaud Condé and Ana Lorena Gutierrez for their critical reading of the manuscript.

