## Supplemental Table and Figures for "The dengue virus NS1 protein alters *Aedes aegypti* midgut permeability and favors virus dissemination"

### Slide 1
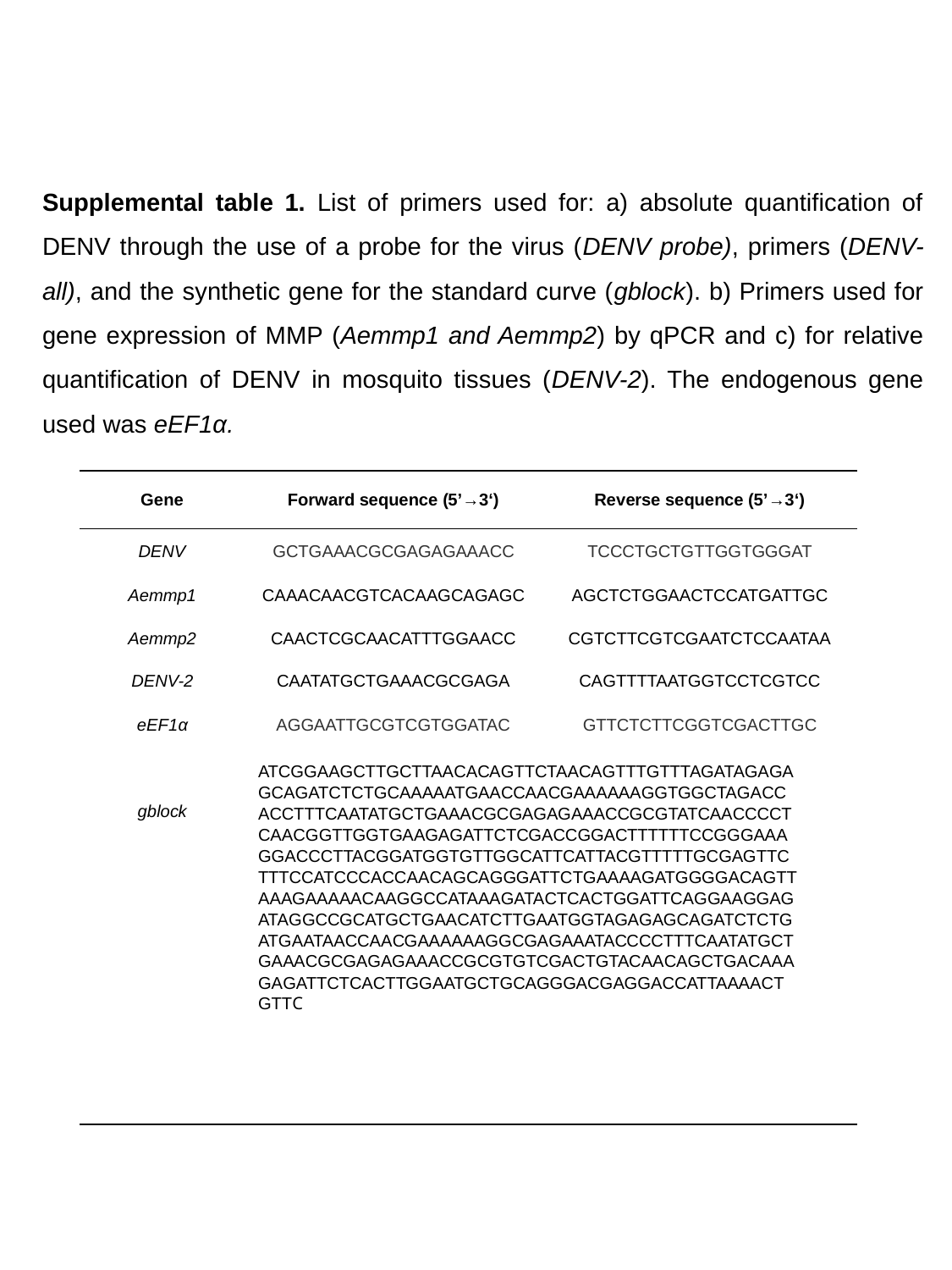

Supplemental table 1. List of primers used for: a) absolute quantification of DENV through the use of a probe for the virus (DENV probe), primers (DENV-all), and the synthetic gene for the standard curve (gblock). b) Primers used for gene expression of MMP (Aemmp1 and Aemmp2) by qPCR and c) for relative quantification of DENV in mosquito tissues (DENV-2). The endogenous gene used was eEF1α.

### Slide 2
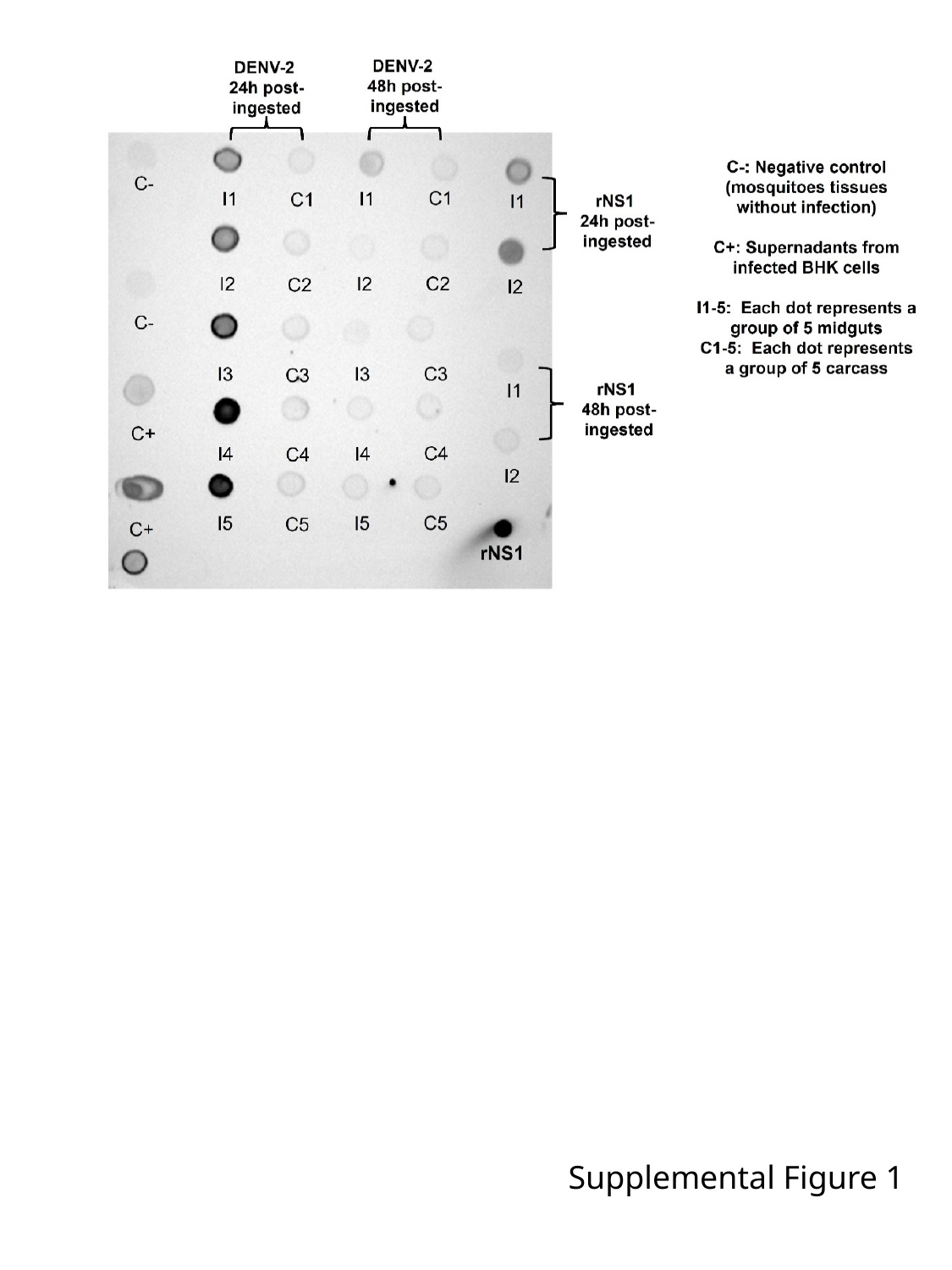

Supplemental Figure 1

### Slide 3
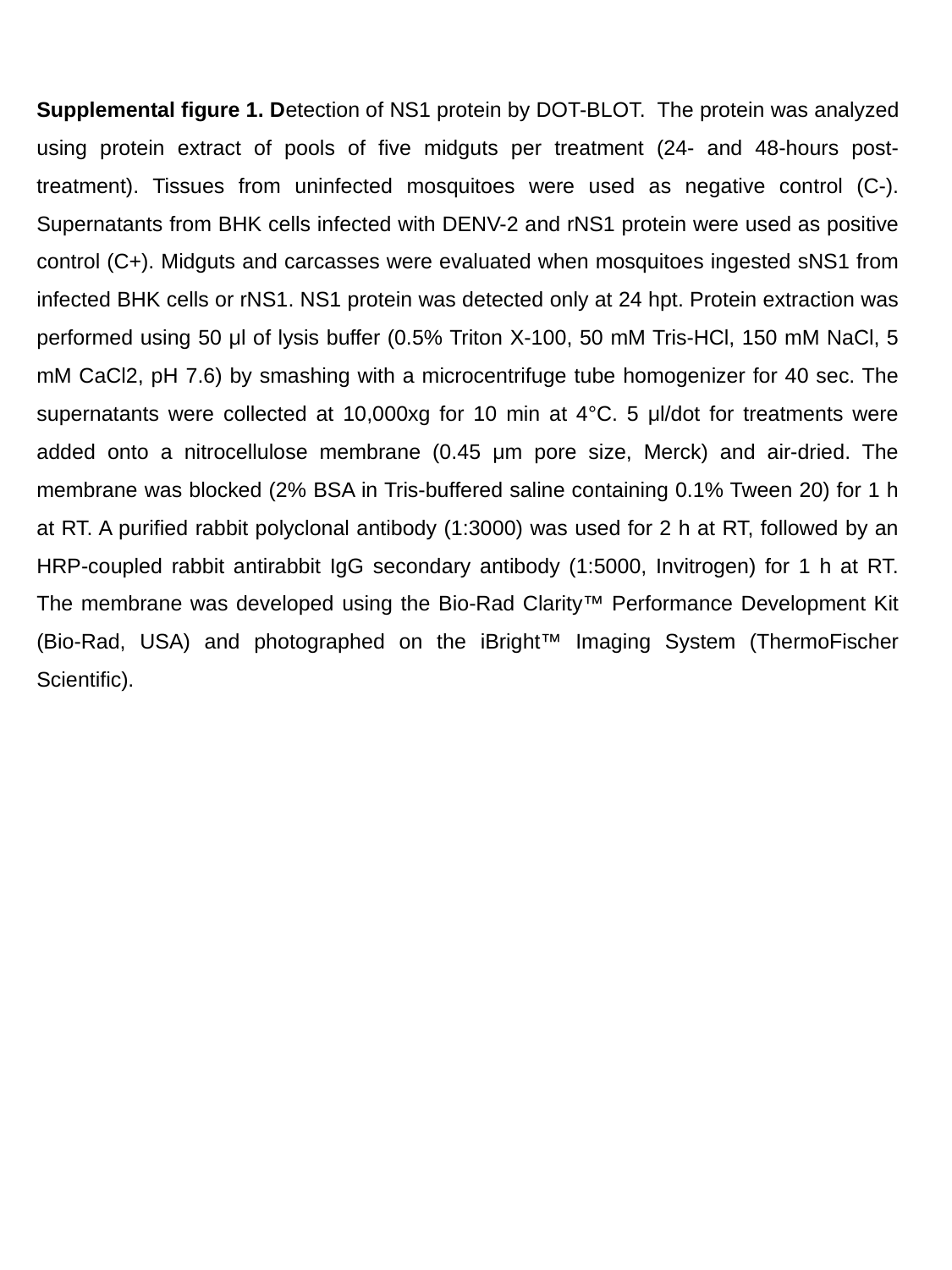

Supplemental figure 1. Detection of NS1 protein by DOT-BLOT. The protein was analyzed using protein extract of pools of five midguts per treatment (24- and 48-hours post-treatment). Tissues from uninfected mosquitoes were used as negative control (C-). Supernatants from BHK cells infected with DENV-2 and rNS1 protein were used as positive control (C+). Midguts and carcasses were evaluated when mosquitoes ingested sNS1 from infected BHK cells or rNS1. NS1 protein was detected only at 24 hpt. Protein extraction was performed using 50 μl of lysis buffer (0.5% Triton X-100, 50 mM Tris-HCl, 150 mM NaCl, 5 mM CaCl2, pH 7.6) by smashing with a microcentrifuge tube homogenizer for 40 sec. The supernatants were collected at 10,000xg for 10 min at 4°C. 5 μl/dot for treatments were added onto a nitrocellulose membrane (0.45 μm pore size, Merck) and air-dried. The membrane was blocked (2% BSA in Tris-buffered saline containing 0.1% Tween 20) for 1 h at RT. A purified rabbit polyclonal antibody (1:3000) was used for 2 h at RT, followed by an HRP-coupled rabbit antirabbit IgG secondary antibody (1:5000, Invitrogen) for 1 h at RT. The membrane was developed using the Bio-Rad Clarity™ Performance Development Kit (Bio-Rad, USA) and photographed on the iBright™ Imaging System (ThermoFischer Scientific).

### Slide 4
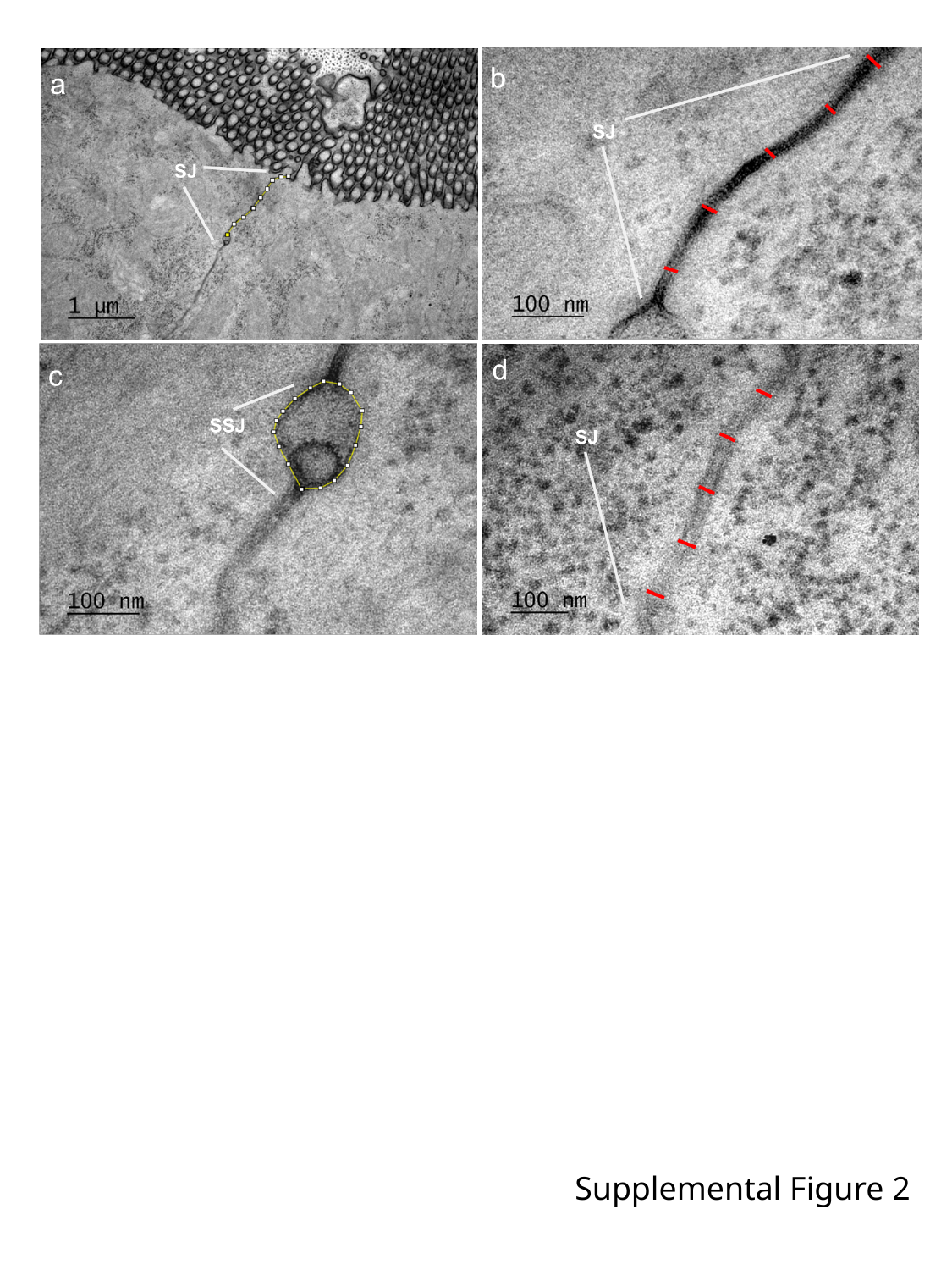

Supplemental Figure 2

### Slide 5
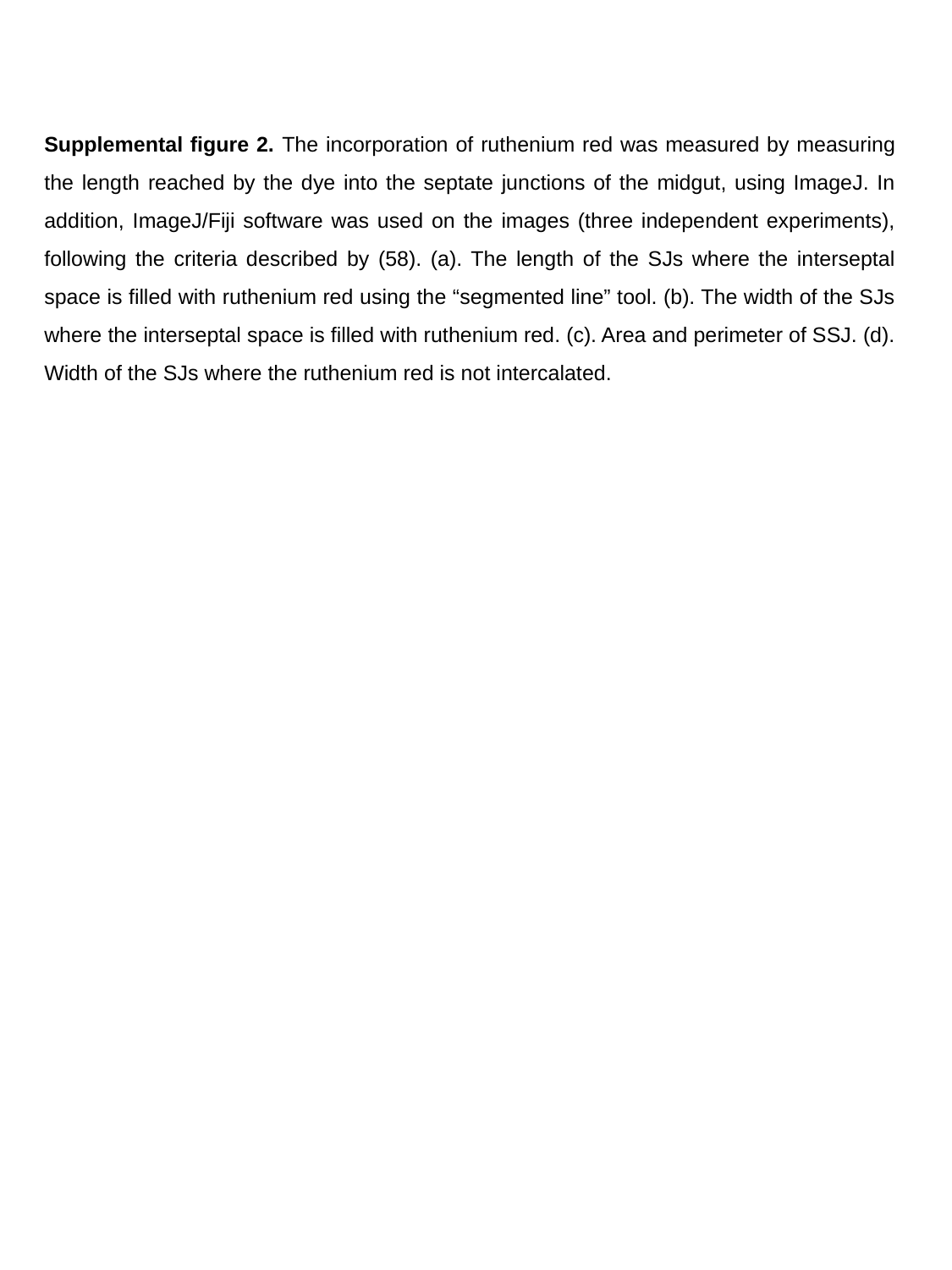

Supplemental figure 2. The incorporation of ruthenium red was measured by measuring the length reached by the dye into the septate junctions of the midgut, using ImageJ. In addition, ImageJ/Fiji software was used on the images (three independent experiments), following the criteria described by (58). (a). The length of the SJs where the interseptal space is filled with ruthenium red using the “segmented line” tool. (b). The width of the SJs where the interseptal space is filled with ruthenium red. (c). Area and perimeter of SSJ. (d). Width of the SJs where the ruthenium red is not intercalated.

### Slide 6
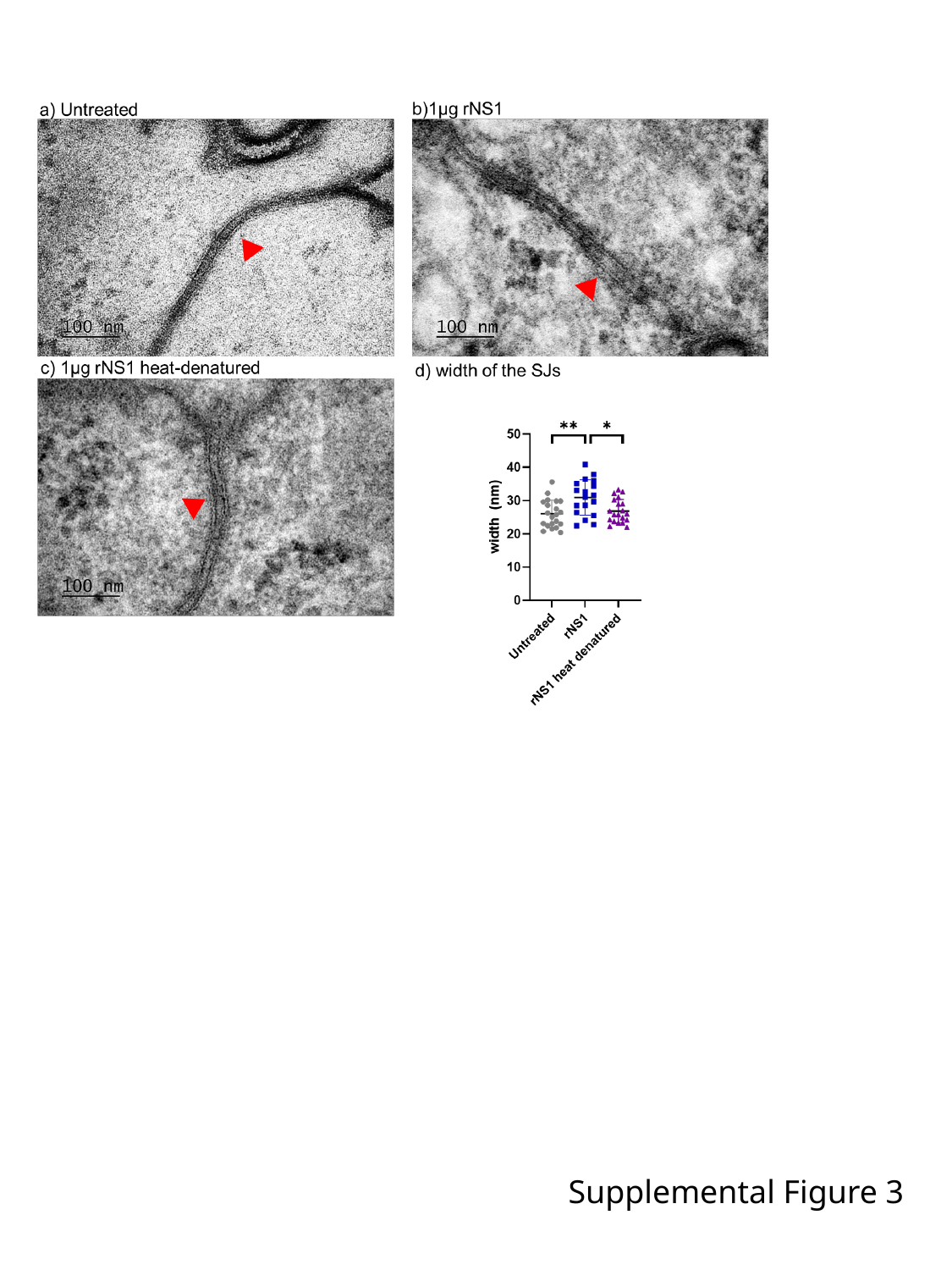

Supplemental Figure 3

### Slide 7
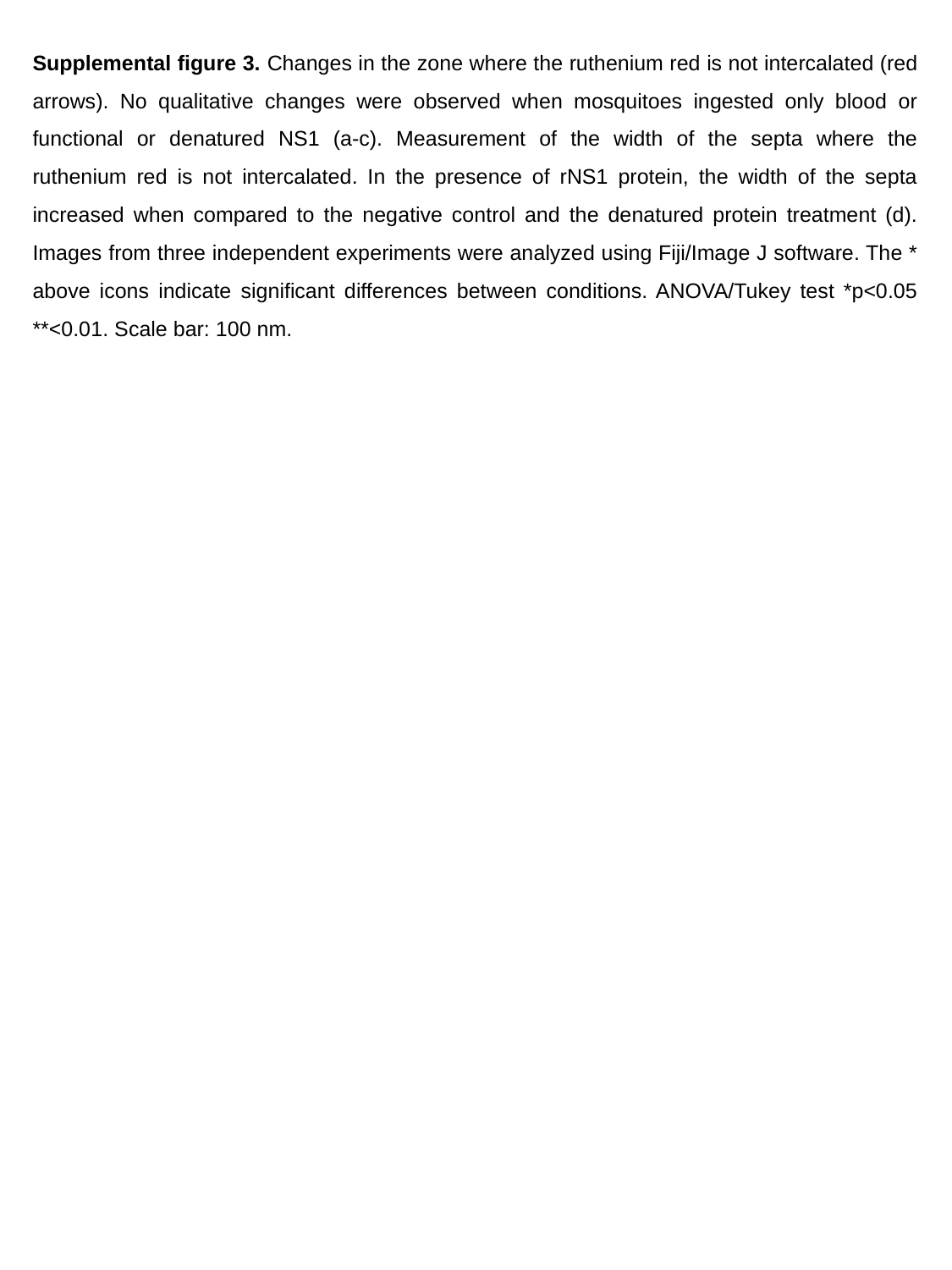

Supplemental figure 3. Changes in the zone where the ruthenium red is not intercalated (red arrows). No qualitative changes were observed when mosquitoes ingested only blood or functional or denatured NS1 (a-c). Measurement of the width of the septa where the ruthenium red is not intercalated. In the presence of rNS1 protein, the width of the septa increased when compared to the negative control and the denatured protein treatment (d). Images from three independent experiments were analyzed using Fiji/Image J software. The * above icons indicate significant differences between conditions. ANOVA/Tukey test *p<0.05 **<0.01. Scale bar: 100 nm.

### Slide 8
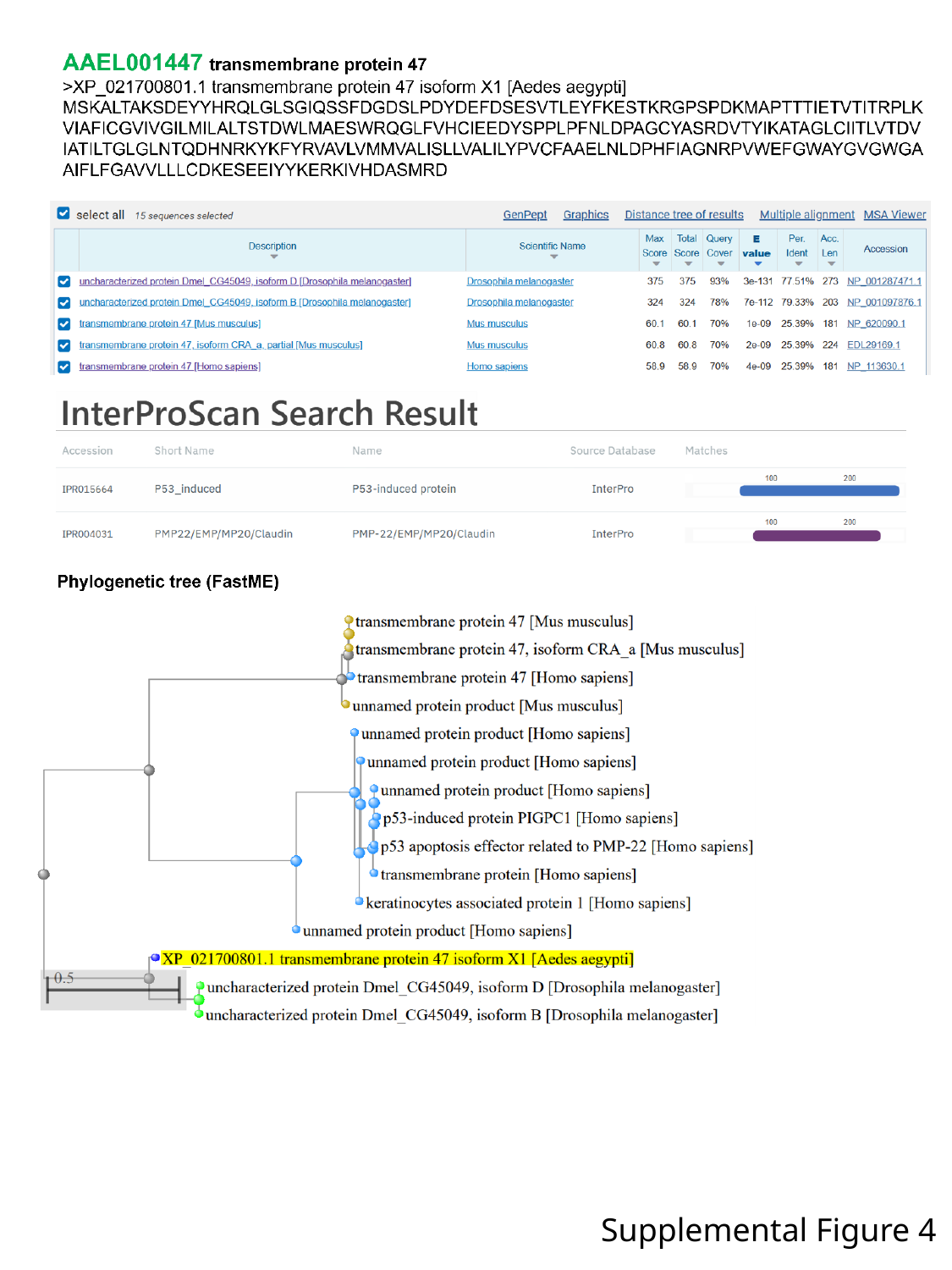

Supplemental Figure 4

### Slide 9
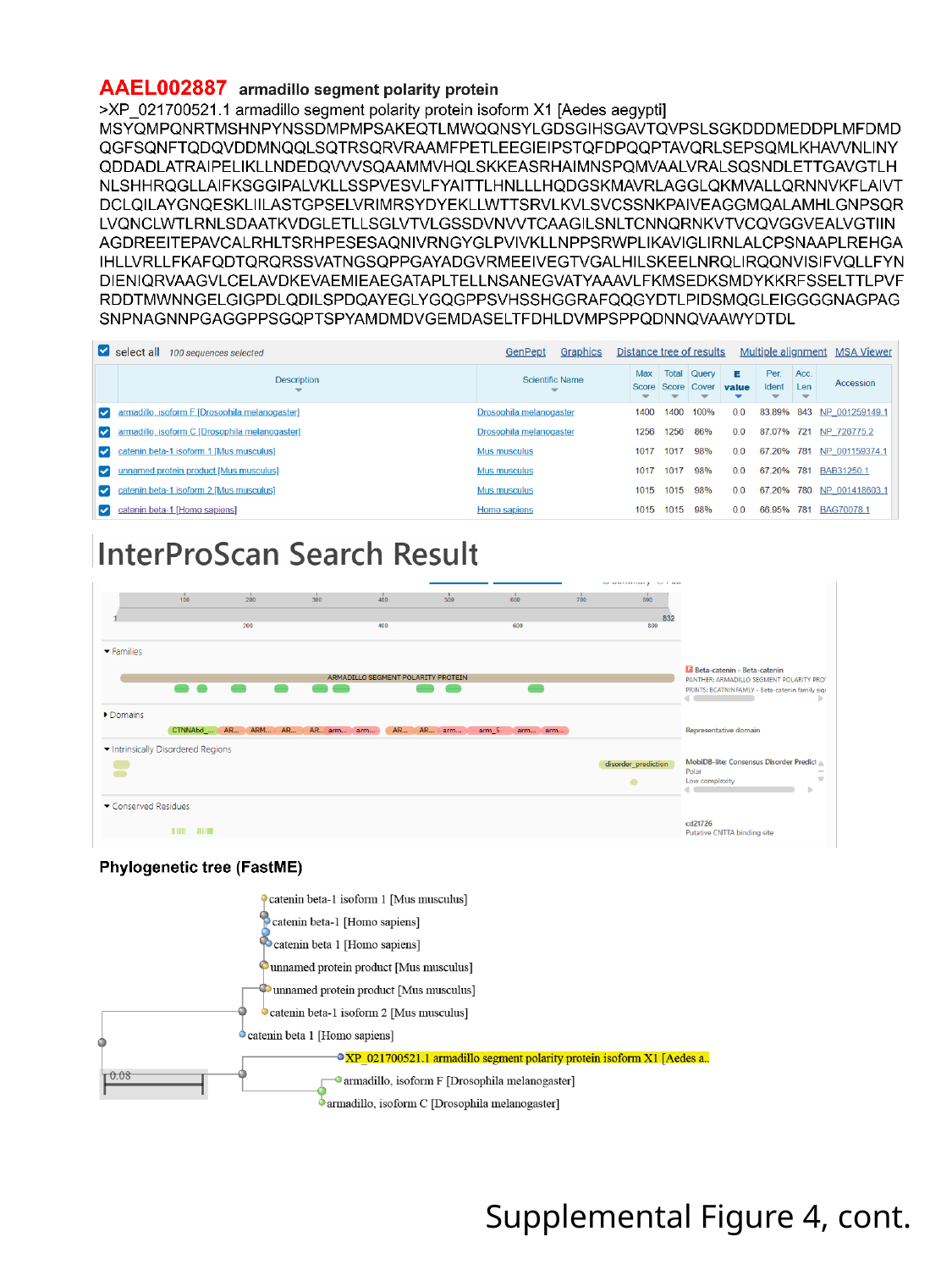

Supplemental Figure 4, cont.

### Slide 10
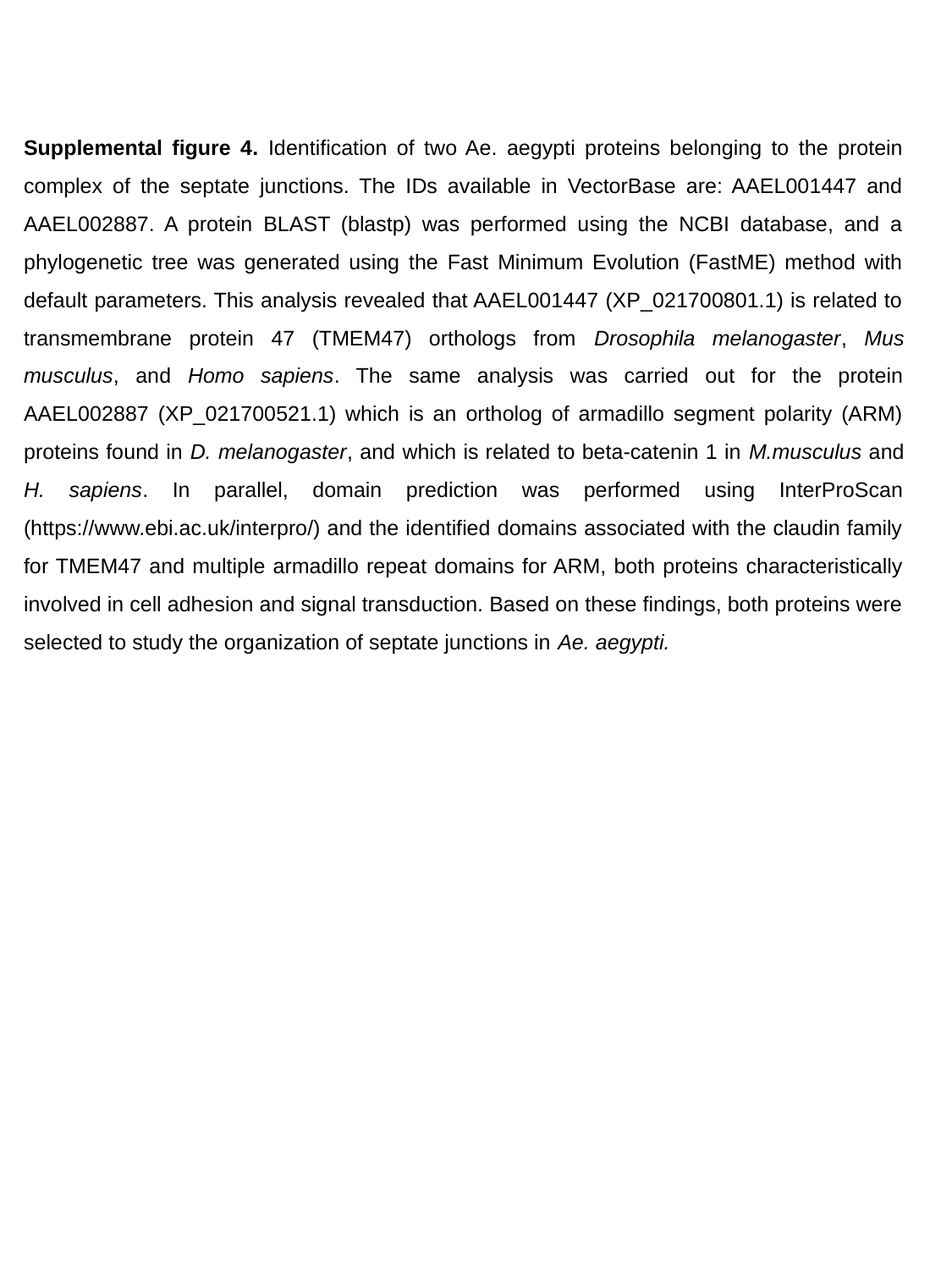

Supplemental figure 4. Identification of two Ae. aegypti proteins belonging to the protein complex of the septate junctions. The IDs available in VectorBase are: AAEL001447 and AAEL002887. A protein BLAST (blastp) was performed using the NCBI database, and a phylogenetic tree was generated using the Fast Minimum Evolution (FastME) method with default parameters. This analysis revealed that AAEL001447 (XP_021700801.1) is related to transmembrane protein 47 (TMEM47) orthologs from Drosophila melanogaster, Mus musculus, and Homo sapiens. The same analysis was carried out for the protein AAEL002887 (XP_021700521.1) which is an ortholog of armadillo segment polarity (ARM) proteins found in D. melanogaster, and which is related to beta-catenin 1 in M.musculus and H. sapiens. In parallel, domain prediction was performed using InterProScan (https://www.ebi.ac.uk/interpro/) and the identified domains associated with the claudin family for TMEM47 and multiple armadillo repeat domains for ARM, both proteins characteristically involved in cell adhesion and signal transduction. Based on these findings, both proteins were selected to study the organization of septate junctions in Ae. aegypti.

### Slide 11
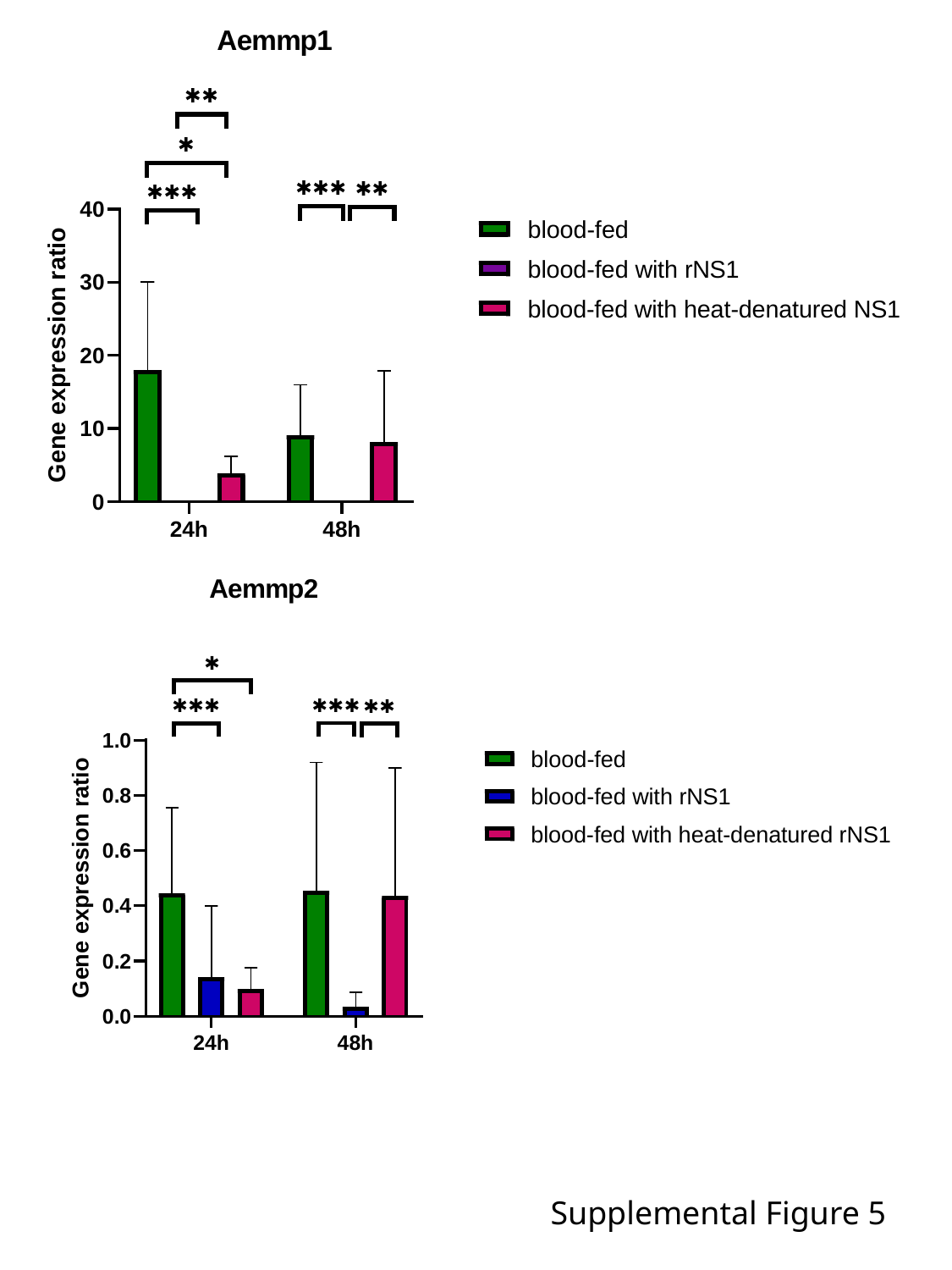

Supplemental Figure 5

### Slide 12
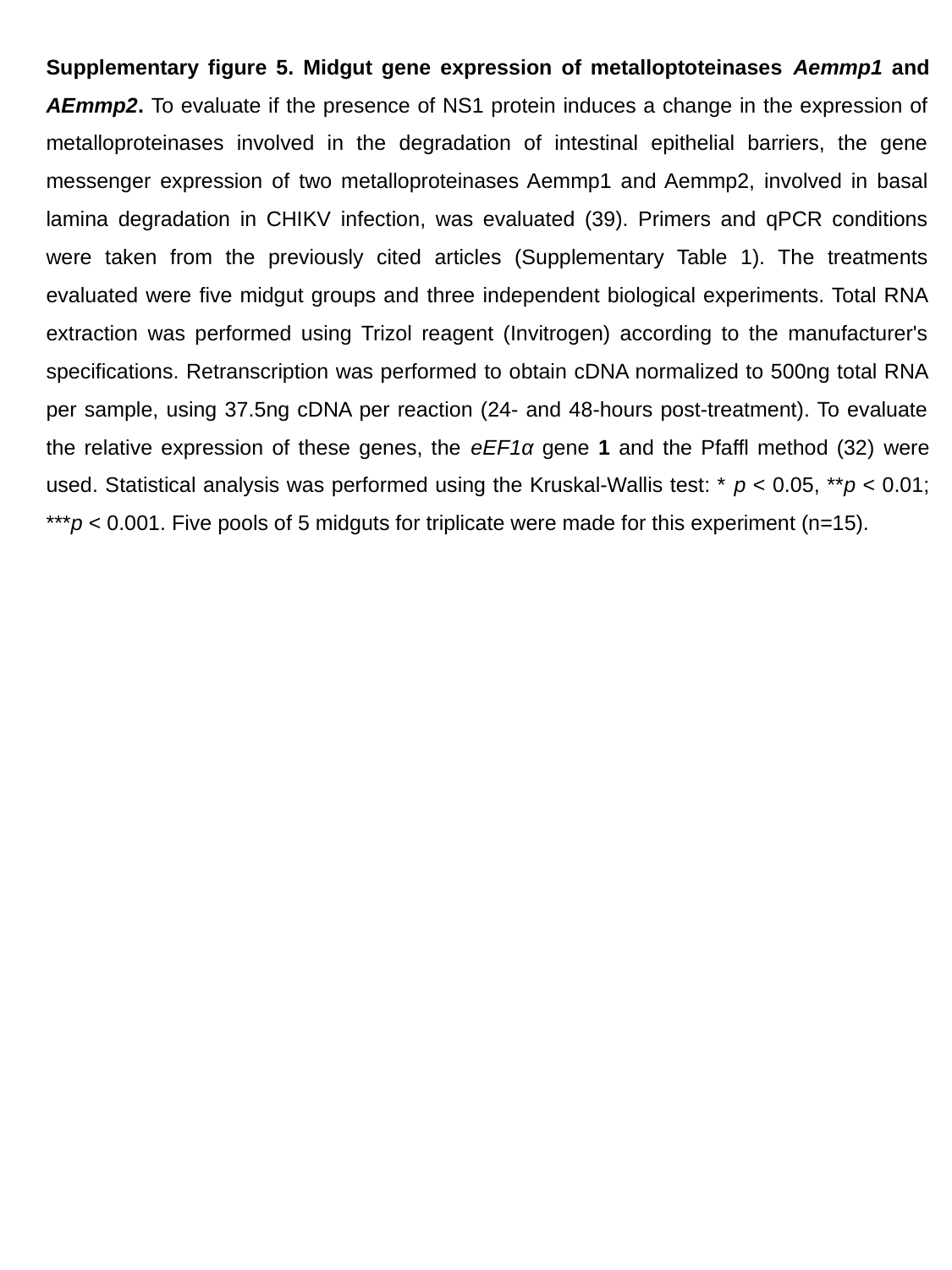

Supplementary figure 5. Midgut gene expression of metalloptoteinases Aemmp1 and AEmmp2. To evaluate if the presence of NS1 protein induces a change in the expression of metalloproteinases involved in the degradation of intestinal epithelial barriers, the gene messenger expression of two metalloproteinases Aemmp1 and Aemmp2, involved in basal lamina degradation in CHIKV infection, was evaluated (39). Primers and qPCR conditions were taken from the previously cited articles (Supplementary Table 1). The treatments evaluated were five midgut groups and three independent biological experiments. Total RNA extraction was performed using Trizol reagent (Invitrogen) according to the manufacturer's specifications. Retranscription was performed to obtain cDNA normalized to 500ng total RNA per sample, using 37.5ng cDNA per reaction (24- and 48-hours post-treatment). To evaluate the relative expression of these genes, the eEF1α gene 1 and the Pfaffl method (32) were used. Statistical analysis was performed using the Kruskal-Wallis test: * p < 0.05, **p < 0.01; ***p < 0.001. Five pools of 5 midguts for triplicate were made for this experiment (n=15).
